# Human milk bacteria assembled into functionally distinct synthetic communities in infant formula differently affect intestinal physiology and microbiota in neonatal mini-piglets

**DOI:** 10.1101/2025.01.21.634066

**Authors:** Charles Le Bras, Gwenaelle Randuineau, Armelle Cahu, Patrice Dahirel, Sylvie Guérin, Regis Janvier, Véronique Romé, Lucie Rault, Marie-Bernadette Maillard, Amandine Bellanger, Yves Le Loir, Sophie Blat, Sergine Even, Isabelle Le Huërou-Luron

## Abstract

The contribution of Human milk (HM) microbiota to infant gut health was addressed by evaluating the impact of HM bacteria, combined in two synthetic communities (SynComs) exhibiting anti-inflammatory (AI) or high immunomodulatory (HI) properties *in vitro*, on gut immune and barrier functions, and microbiota. Neonatal mini-piglets were fed either a formula without supplementation (CTRL) or supplemented with AI or HI SynComs, and were compared to sow milk-fed (SM) piglets over a period of 24 days. Feces were collected on postnatal day (PND) 8, and ileal, colonic and fecal samples were collected on PND24. The multifactorial analysis indicated that the two HM-derived SynComs impacted microbiota and intestinal functions differently. Several genera, mainly belonging to Bacillota, displayed different relative abundances between the formula-fed groups at both PND8 and PND24. At PND8, the fecal sIgA level in HI piglets was slightly lower than in SM piglets but markedly higher than in CTRL and AI piglets. SynComs HI and/or AI slightly increased the expression of genes involved in pro-inflammatory (IL6, TNFaR1), antioxidant (SOD2), anti-inflammatory (SOCS3) and Treg (FOXP3) pathways in ileal and colonic tissues compared with the CTRL group. Systemic immune functions were also modulated with a cytokine secretion capacity of peripheral blood mononuclear cells that tended to be higher with HI supplementation. Interestingly, SynCom bacteria were correlated with several ileal and colonic genera, and both were correlated with physiological variables. Overall, our findings support the influence of HM bacteria, provided in formulas as SynCom at a physiological concentration, on gut microbiota and functions.

**Importance:** Early-life environmental factors, such as neonatal diet, influence the gut microbiota, which plays a key role in the functional development of the gut. However, the role of the human milk (HM) microbiota, particularly with regard to the immunomodulatory properties of HM bacteria, is not well understood. This study investigates the differential effects of two synthetic communities with a similar taxonomic composition representative of the taxonomic diversity of the HM microbiota. Thse communities exhibit contrasting immunomodulatory properties that were previously characterized using an *in vitro* intestinal quadricellular model. Daily supplementation with these two SynComs modulated the composition of the gut microbiota and the gut physiology differently, particularly the intestinal immune signatures. In conclusion, the functional profile of bacteria within the HM microbiota may induce distinct developmental profiles of gut physiology in infants.

## Introduction

Optimal infant growth and development, particularly with regard to gut digestive and immunological functions and microbiota, depend on early nutrition (1). Human milk (HM) is the optimal source of nutrition for infants in their first few months of life (2, 3). The beneficial properties of HM are mediated by bioactive compounds that promote the maturation of the gut immune system in interaction with the development of the gut microbiota (4, 5). Breast milk contains a diverse bacterial community, with over 200 different species belonging to dozens of genera identified (6, 7). That includes the prevalent genera *Bifidobacterium* and *Lactobacillus*, as well as *Streptococcus, Staphylococcus, Cutibacterium* and *Corynebacterium* (6, 8, 9). Despite the low bacterial load in HM (10^3^-10^4^ colony-forming units (CFU)/mL), these bacteria contribute to the neonatal gut microbiota, accounting for 5-33 % of the total bacteria present in the feces of breastfed infants under six months of age (10, 11). Beyond their direct role in seeding the infant microbiota, HM bacteria are likely to contribute to the development of the gut microbiota. This is achieved by competing with gut microbes for nutrients or mucosal binding sites, directly inhibiting them, or contributing to trophic chains (4). HM bacteria may also influence the development of the gut immune system. This occurs via their immunomodulatory properties, such as modulation of cytokine production and the induction of secretory IgA (sIgA), or via their impact on gut barrier function (12). The immunomodulatory activity of HM bacteria may be associated with the presence of bacterial surface antigens or intracellular antigens, released during digestion, as well as their metabolism. Some HM bacteria can digest human milk oligosaccharides (HMOs) and other milk compounds, producing short chain fatty acids (SCFAs) with significant immunomodulatory capacities (7). Studies *in vitro* and *in vivo* in mice and piglets have shown that HM bacteria belonging to the genera *Bifidobacterium*, *Lactobacillus* and *Streptococcus* exhibit a wide variety of immunomodulatory profiles (13–17). With the exception of these genera, little information is available on the immunomodulatory properties of HM strains belonging to other genera. However, bacteria belonging to these genera that have been isolated from other types of microbiota, such as skin, mouth or gut microbiota, have also demonstrated strain-dependent immunomodulatory capacities, and impacted barrier function (18–21). This suggests that HM-derived bacteria can impact gut physiological functions. Nevertheless, the impact of HM-derived bacteria assembled in complex communities that mimic the actual bacterial communities present in HM, at a low but continuous dose, on gut functions and microbiota has been insufficiently addressed so far.

To further explore the *in vivo* role of HM bacteria assembled in synthetic bacterial communities (SynComs) on intestinal homeostasis, Yucatan piglets were fed dairy-based infant formula supplemented with HM SynComs. The suckling pig is a well-established and suitable animal model for human infants, with a digestive system that is well characterised and closely resembles that of humans (22–25). Furthermore, Yucatan piglets can consume a diet consisting entirely of dairy-based infant formula and provide sufficient tissue and digesta samples for various analyses (26). A sequential microbial colonization of the digestive system with a progressive increase in α-diversity occurs during suckling and especially at weaning in the piglet as in the human infant (27–29). Two SynComs of 11 HM strains each, closely related in terms of taxonomy and covering the prevalent taxa of HM microbiota were used. We have previously demonstrated *in vitro* that the intestinal epithelium was differently affected depending on the HM strain assembly, these two SynComs displaying an anti-inflammatory profile (AI) *vs.* a high immunostimulatory (HI) one (9). It was hypothesized that these two HM SynComs added to infant formula at a concentration close to physiological concentrations (∼5.5.10^5^ CFU/mL infant formula) would differentially affect key gut immune and barrier functions *in vivo*, according to their functional characteristics, in association with changes in gut microbiota composition. The gut immune and barrier functions and microbiota were also examined in sow milk-fed (SM) piglets as a natural physiological control group.

## Results

### SynCom supplementation of formulas was well tolerated and did not affect piglet growth

From postnatal day (PND) 2 to PND24, Yucatan piglets were fed dairy-based infant formula (Supplemental Table S1) without supplementation (CTRL) or supplemented with SynComs AI or HI (Table 1; (9)). The AI and HI bacterial doses added in formulas corresponded to 5.5 x 10^5^ CFU/mL formula, amounting to an average daily intake of 1.28 x 10^8^ CFU/kg body weight (BW) for piglets fed supplemented formulas. During the whole experimental period, formula-fed piglets remained healthy, consumed an average value of 255 ± 3 mL/kg BW, with an average daily BW gain of 74 ± 14 g/day (Figure 1). Supplementing formulas with SynComs had no significant effect on BW gain compared to the CTRL, though a significant sex effect was observed (P=0.045) (Figure 1). As expected, SM piglets showed significantly higher growth (P<0.05) with an average daily BW gain of 146 ± 4 g/day compared to the formula-fed piglets.

**Figure 1.**
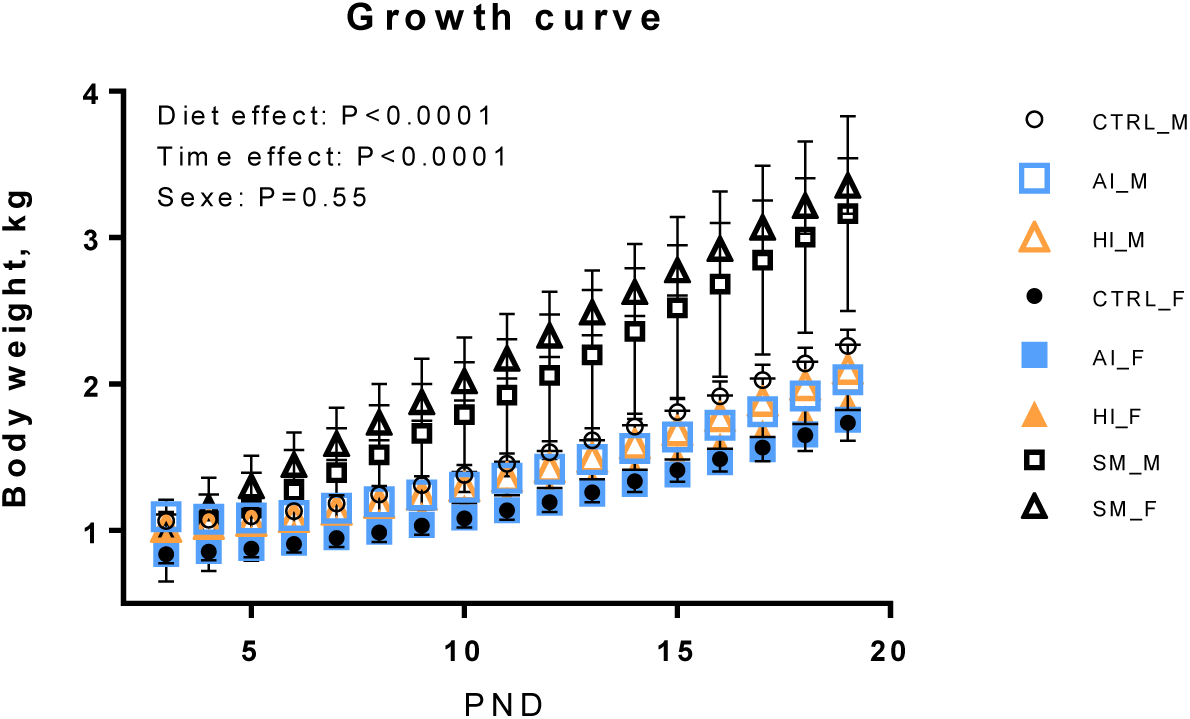
Growth curve of female (F) and male (M) formula-fed piglets (CRTL, AI, HI) and sow milk-fed (SM) piglets. Data are shown as means ± SEM. Body weight differences between the CTRL, AI, HI and SM groups were assessed using repeated-measures ANOVA, testing for the effects of diet, time and sex. The body weight of CTRL, AI and HI piglets differed from that of SM piglets from PND7 to PND19 (p < 0.05).

**Table 1.**
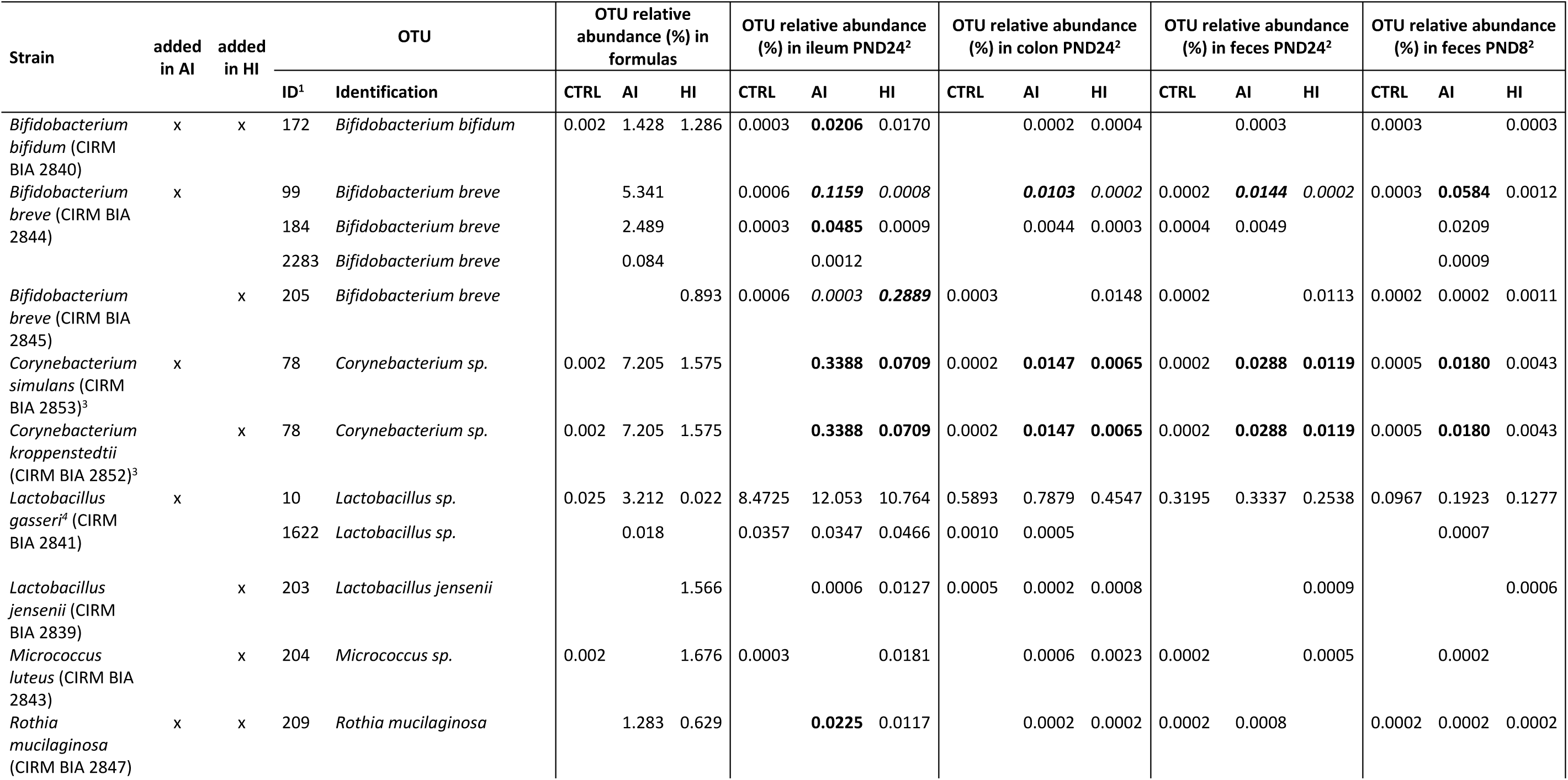
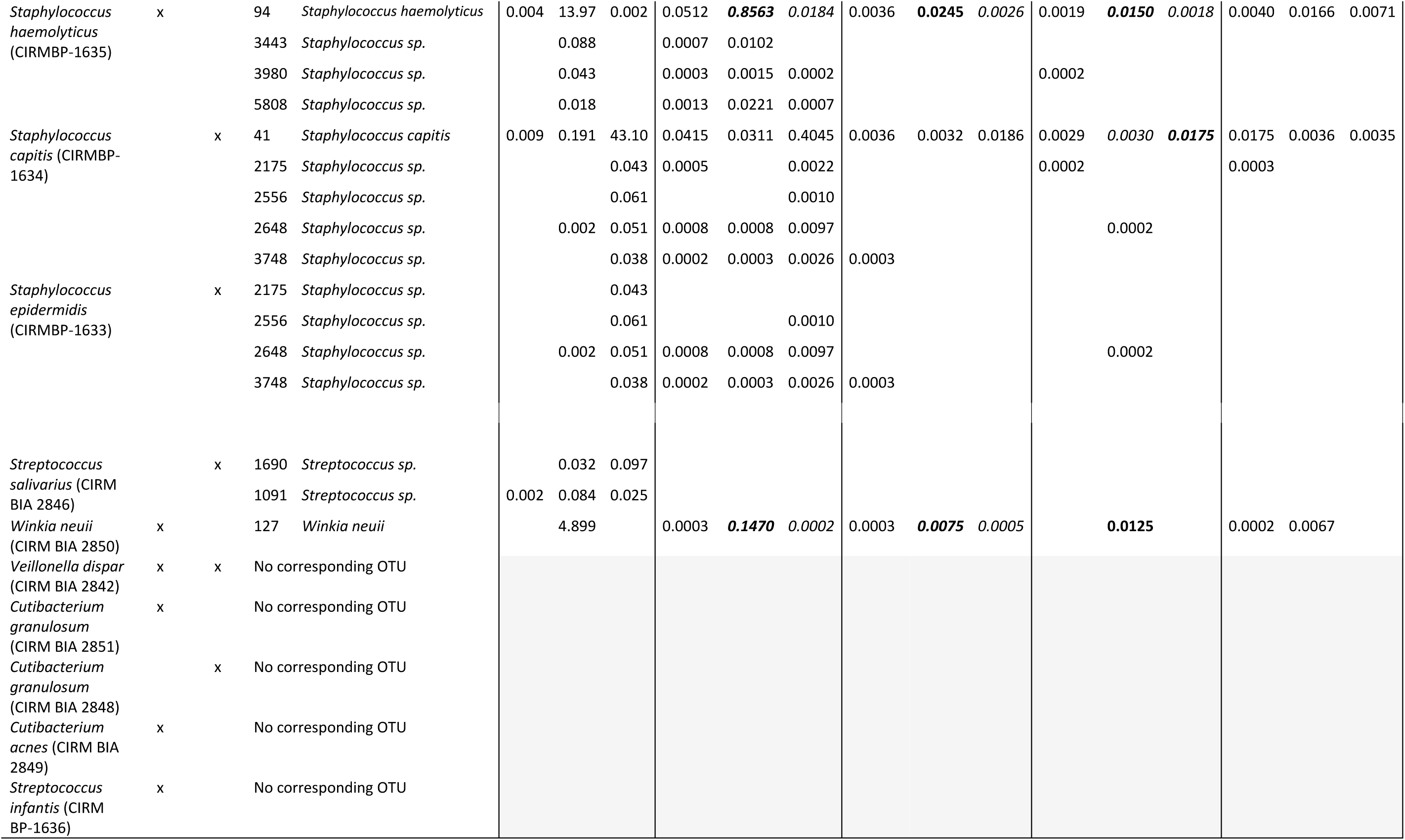
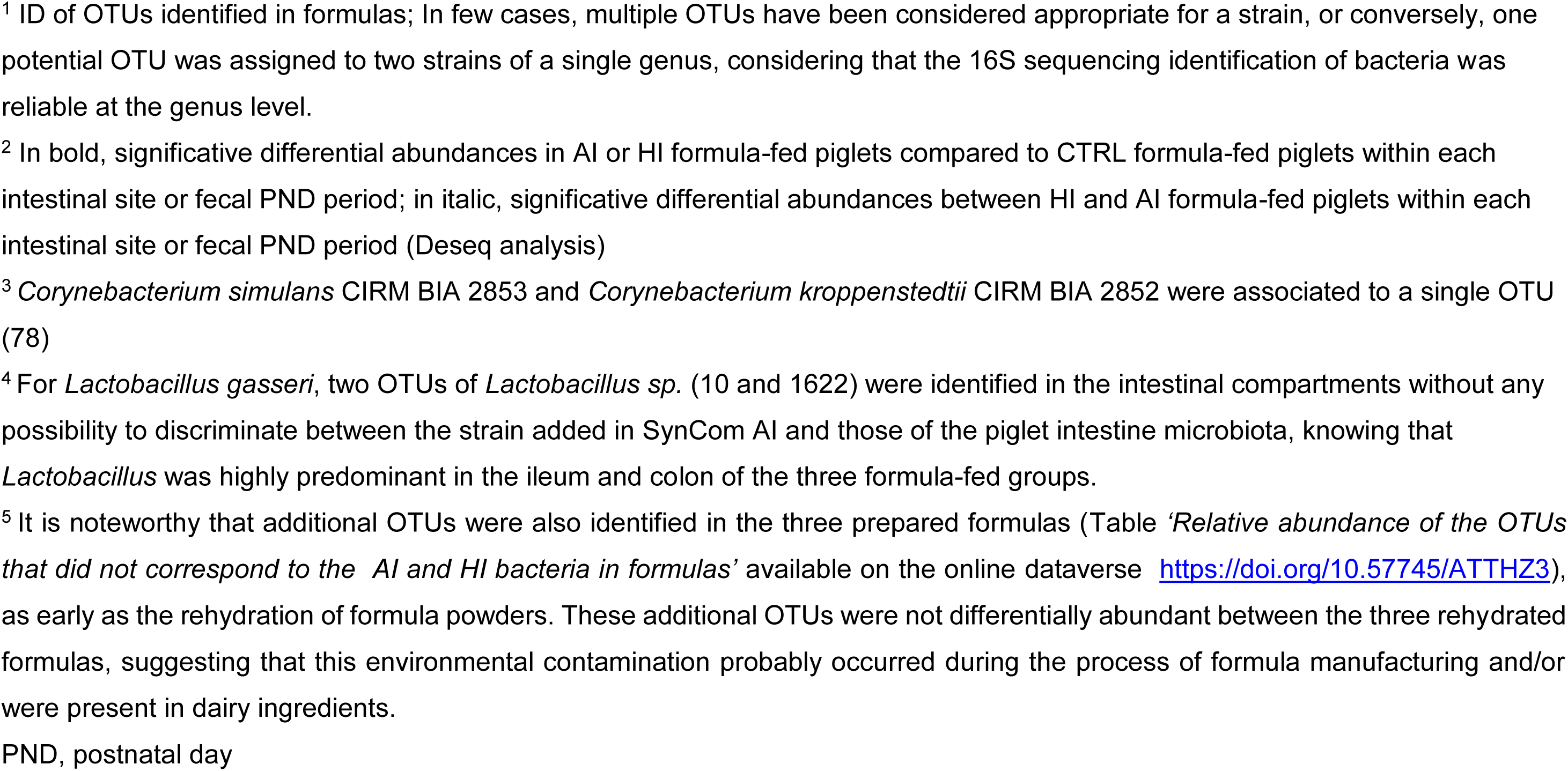
Relative abundances of the Syncom OTUs in formulas^5^, and in the feces at PND8 and ileum, colon and feces of formula-fed (CRTL, AI and HI) piglets at PND24. OTUs potentially corresponding to the strains added in SynCom AI and/or HI formulas are presented together with their relative abundance (in % of total OTUs) in feces, ileon and colon microbiota.

### Overall impact of SynCom AI and HI on the intestinal homeostasis of formula-fed piglets

A multifactorial analysis (MFA) was conducted in order to analyze the overall effect of SynCom supplementation of formulas on intestinal homeostasis. To this purpose, Partial Least Squares-Discriminant Analysis (PLS-DA) was first performed on the variables related to microbiota composition, SCFA concentration, Ussing chamber permeability, histomorphometric data, sIgA content and gene expression measured in the ileum and colon as well as cytokine and sIgA production of ileal Peyer Patch (PP) cell culture, from the three formula-fed piglet groups (dataset accessible in the repository https://doi.org/10.57745/ATTHZ3) . The most discriminant variables with a Variable Importance in Projection (VIP) score >1 in the PLS-DA (n = 112 out of a total of 201 variables in the ileum and n = 141 out of a total of 250 variables in the colon) were selected for the MFA after grouping by function (barrier, immunity, microbiota activity, nutrient transporter, endocrine and tryptophan metabolism) or phylum (Actinomycetota, Bacillota, Bacteroidota, Pseudomonadota, and ’Other Phyla’) as detailed in Materials & Methods section (Supplemental Figure S1, dataset accessible in the repository https://doi.org/10.57745/ATTHZ3). MFA was able to distinguish the three groups of formula-fed piglets in the ileum and in the colon (Figure 2). The groups of variables contributing to the discrimination between formula-fed piglet groups included immunity, microbiota (Actinomycetota, Bacillota and Pseudomonadota phyla) and endocrine functions in both the ileum and colon, as well as tryptophan metabolism and barrier functions in the ileum only.

**Figure 2.**
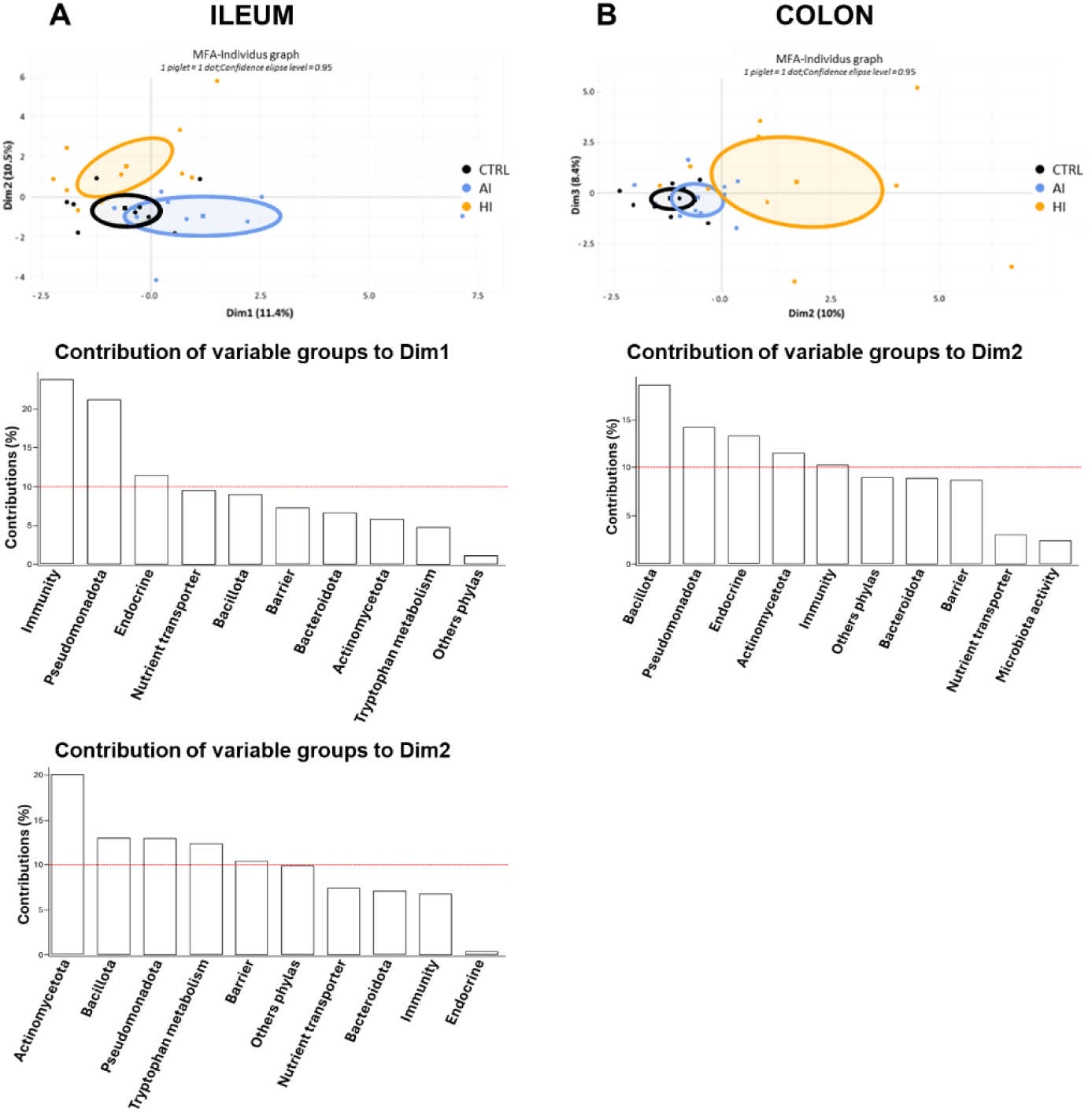
Ileal (A) and colon (B) multifactorial analysis (MFA) using microbiota and intestinal variables of formula-fed piglets (CRTL, AI, HI), and grouped by function or phylum (n = 11 groups of quantitative variables). Variables used in the MFA were the most discriminating variables preselected by Partial Least Squares-Discriminant Analysis (PLS-DA) with a Variable Importance in Projection (VIP) score > 1 for components 1 and 2 (n=112 and 141 variables in the ileum and colon, respectively; Supplemental Figure S1 and Table ‘*Table Individual ileal and colonic data used for PLS-DA and MFA*’ accessible in the repository https://doi.org/10.57745/ATTHZ3). On the MFA plots, the dots represent the piglet individus, and the square symbols the barycenter of the ellipses. The Red dot line on the histograms of contribution of variable groups represents the level of statistical significance corresponding to the inverse of the number of groups of variables used in the analyses.

### SynCom bacterial strains were present in AI and HI intestinal microbiota

To monitor the presence of SynCom bacteria in the intestine of piglets, metabarcoding was applied to AI- and HI-supplemented formulas in order to identify the OTU potentially corresponding to each SynCom bacteria (hereafter referred as to SynCom AI/HI OTUs) in supplemented formulas and in the ileal and colonic piglet digesta. OTUs corresponding to the genera added in SynComs AI and/or HI were identified in AI and/or HI formulas, with high homology between the OTU sequence and the 16S sequence of the SynCom strains, at the exception of *Cutibacterium*, *Veillonella* and a *Streptococcus infantis* strain present in SynCom AI (Table 1, Supplemental Table S2). The absence of corresponding OTUs for *Cutibacterium* and *Veillonella* strains was likely due to the failure of V3-V4 amplification on these strains with the primer pair used, as confirmed by the failure of virtual PCR using the NCBI PrimerBLAST tool. Furthermore, a bias in the lysis of bacterial cells for certain strains that may be resistant to the method used, may have limited DNA recovery from these strains, contributing to their non-detection in formulas or piglet microbiota. Six SynCom AI OTUs (potentially corresponding to the *Bifidobacterium bifidum, Rothia mucilaginosa, Corynebacterium simulans, Bifidobacterium breve, Staphylococcus haemolyticus* and *Winkia neuii* strains added in AI SynCom) exhibited significant higher relative abundances in the ileal microbiota of AI piglets compared to that of CTRL piglets (Figure 3, Table 1). Likewise, the relative abundance of three SynCom HI OTUs (potentially corresponding to the *Corynebacterium kroppenstedtii, Bifidobacterium breve* and *Staphylococcus capitis strains* added in HI) was significantly higher in the ileal microbiota of HI compared to CTRL. Three additional SynCom HI OTUs (potentially corresponding to *Staphylococcus epidermidis, Micrococcus luteus* and *Lactobacillus jensenii*) exhibited a similar higher abundance profile, yet this was not statistically significant. The presence of SynCom OTUs was also observed in the colonic and fecal microbiota of supplemented piglets at PND8 and PND24, yet to a lesser extent, probably due to the very low abundance of these OTUs in these compartments. When found, the relative abundance of SynCom OTUs can represent from 0.01% to 0.86% of total OTUs in ileum, from 0.0002% to 0.02% of total OTUs in colon, and from 0.0002% to 0.06% of total OTUs in feces. Overall, these results suggest that most SynCom bacteria were recovered from the ileal microbiota and, to a lesser extent, the colonic microbiota of supplemented piglets.

**Figure 3.**
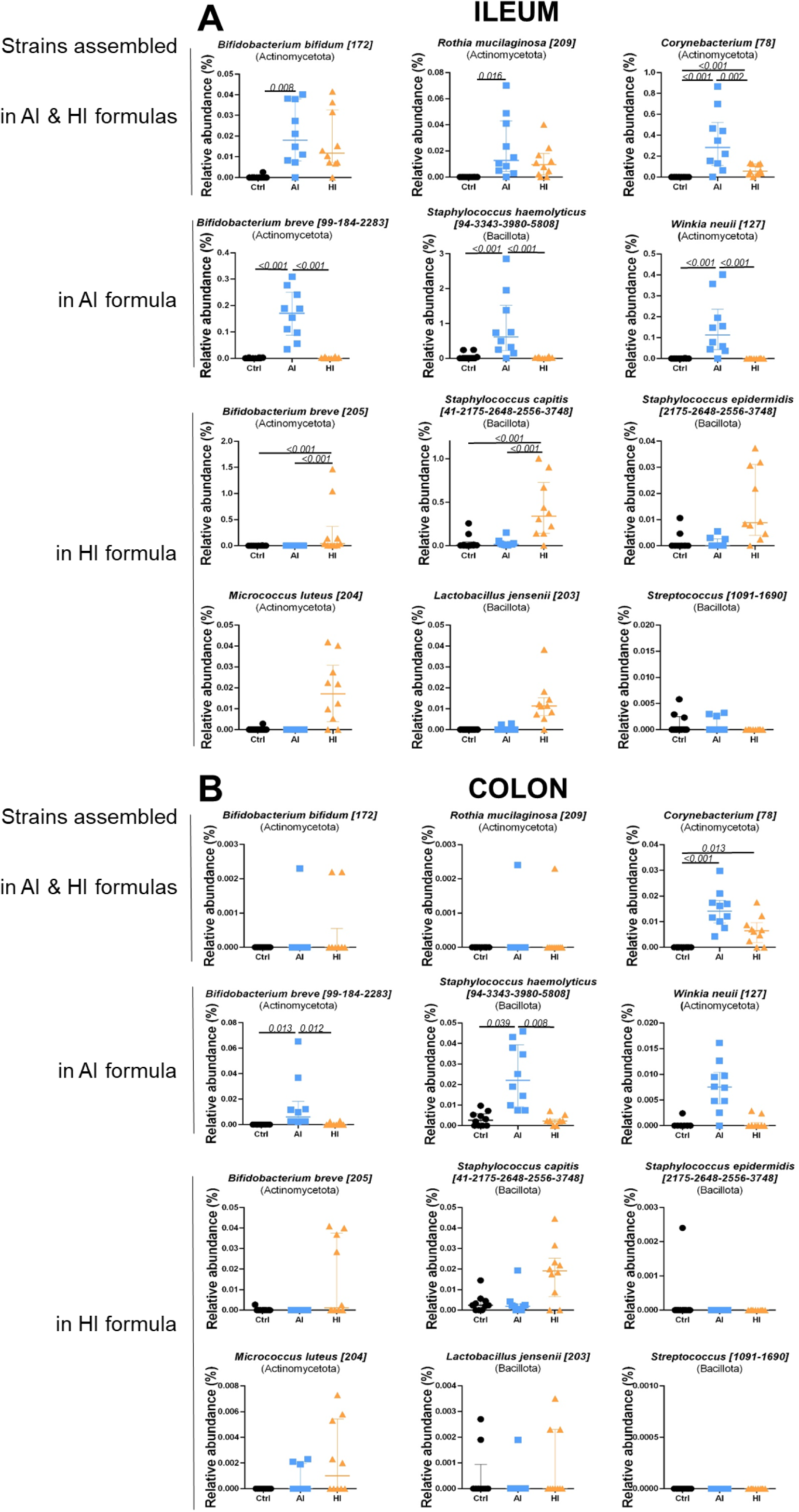
Ileal (A) and colonic (B) relative abundances of the OTUs potentially corresponding to the SynCom strains added in AI and/or HI formulas. Each individual is represented with one dot; Median with interquartile range. A three-way ANOVA was performed testing for main effects of diet, sex, litter, and for interactions between diet and sex, and diet and litter, followed by Tukey *post hoc* test for parametric variables or a Kruskal Wallis test associated with Dunn test for non-parametric variables.

### SynCom supplementation modified microbiota composition and activity

Microbiota was analyzed on feces from formula-fed and SM-fed piglets at PND8 to detect early dietary effects and on ileal and colonic digesta and feces at PND24. Moderate differences were observed in ileal, colonic and fecal microbiota of formula-fed piglets (CTRL, AI, HI). The ileal α-diversity was lower in CTRL than in HI, AI group value being intermediate (Supplemental Figure S2). β-diversity did not differed between formula-fed groups (Supplemental Figure S2). However, consistent with the MFA results, several taxa showed different relative abundances in the ileal and colonic microbiota of the formula-fed groups that go beyond the presence of the SynCom OTUs (Table 2; Supplemental Table S3). At PND24, several families (Lachnospiraceae, Peptostreptococaceae, Ruminococaceae and Erysipelotrichaceae) within Bacillota were more abundant in the ileum of HI compared to CTRL and AI. An intermediate situation was sometimes observed for AI, as for Peptostreptococcaceae, whose relative abundance in AI was not significantly different from HI or CTRL. Some changes were common to the AI and HI groups compared to the CTRL group, notably a higher abundance of Actinomycetaceae and *Actinomyces*.

**Table 2.**
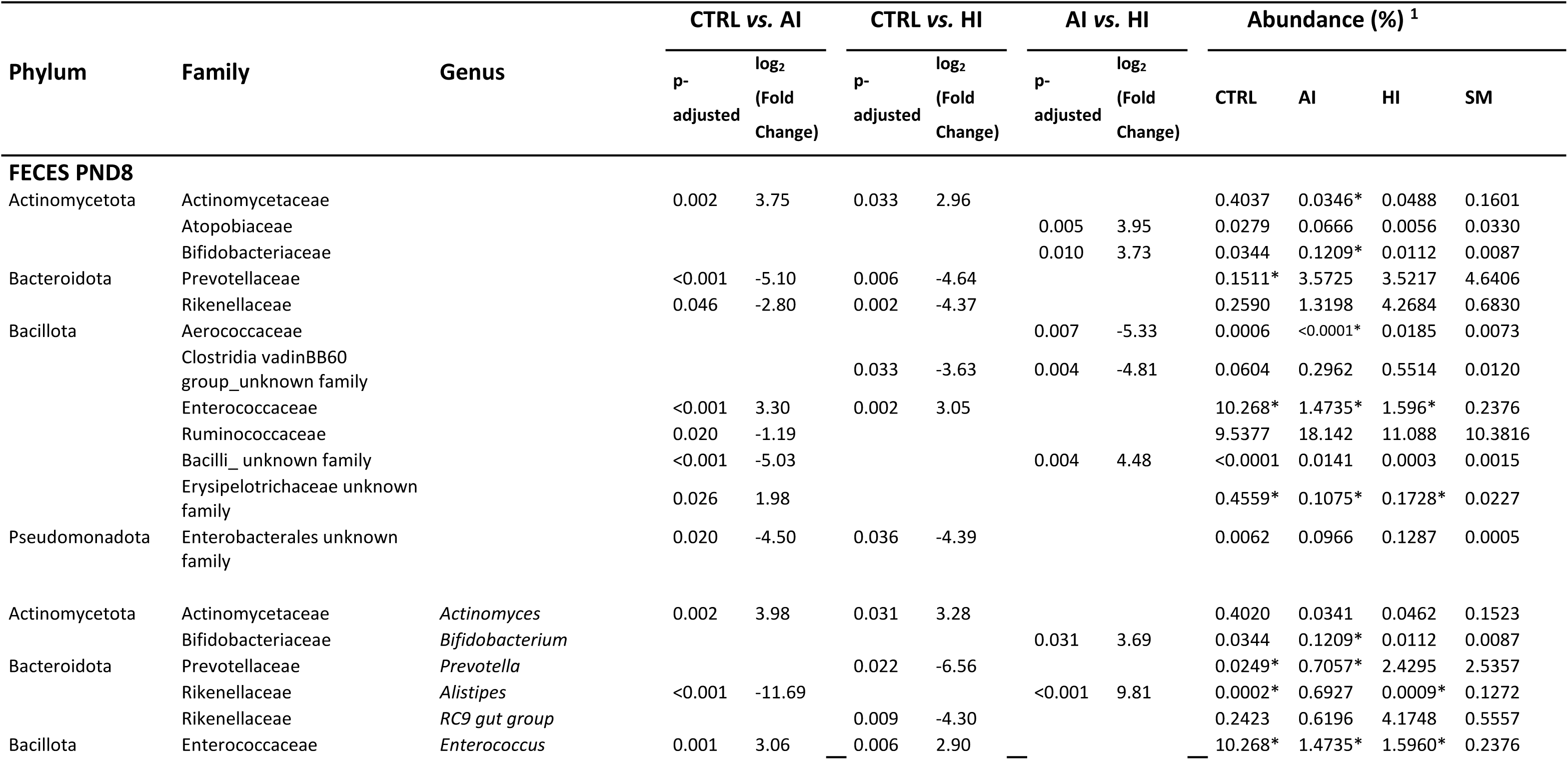
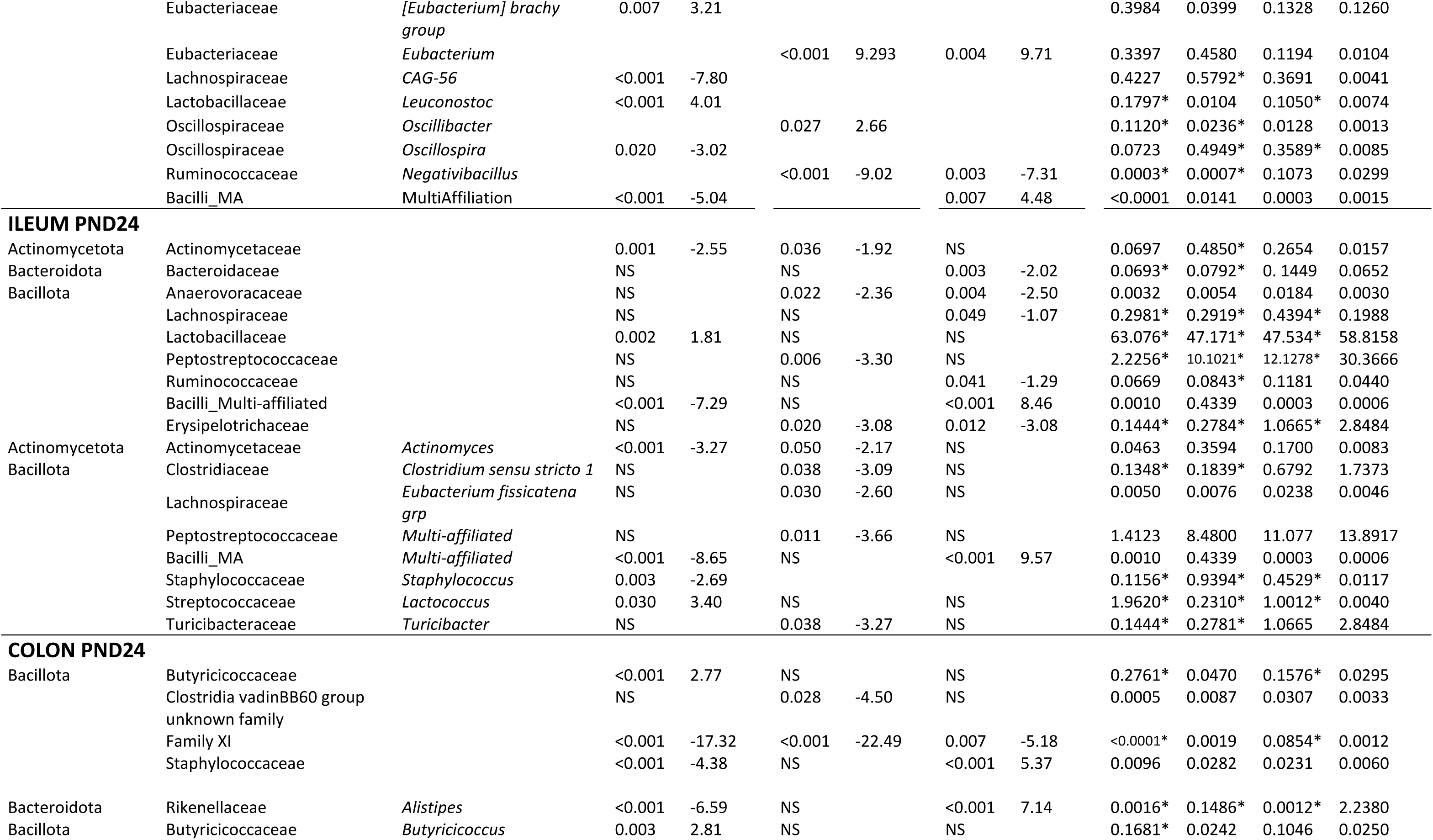
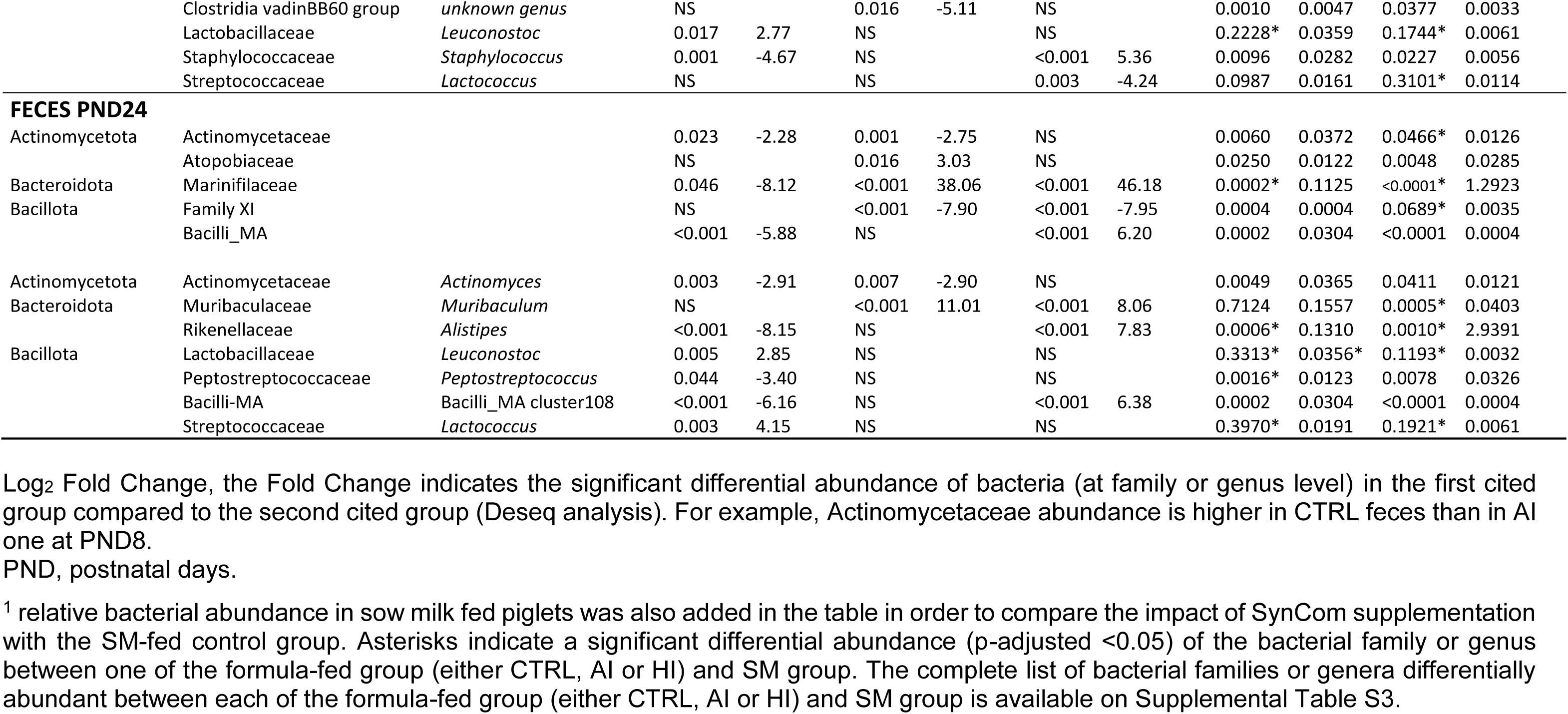
Impact of SynComs AI and HI on the relative abundance of bacterial families, and genera in the feces at PND8 and in the ileum, colon and feces at PND24 of formula-fed and sow milk-fed piglets, and the differential abundance between groups (log_2_ fold change).

In the colon, less differences were observed between formula-fed groups. Family IX and Clostridia vadinBB60 group were more abundant in HI (and, to a lesser extent in AI), while *Allistipes* and *Staphylococcus* were higher in AI. Some ileal and colonic differences were reflected in fecal microbiota at PND24, with higher levels of family XI, Actinomycetaceae, *Actinomyces* and *Allistipes* in AI and/or HI groups compared to CTRL group (Table 2).

At PND8, several taxa showed differential abundance between groups. Differences included a higher abundance of Prevotellaceae/*Prevotella* and Rikenellaceae in AI and HI, Bifidobacteriaceae/*Bifidobacterium* and Ruminococcaceae specifically in AI, and Aerococcaceae and an unknown family within Clostridia vadinBB60 group in HI. Conversely, Actinomycetaceae/*Actinomyces* and Enterococcaceae/*Enterococccus* were less abundant in Syncom-supplemented groups.

The ileal, colonic and fecal microbiota composition of the formula-fed groups was compared with that of SM piglets. Of note, the sow milk microbiota was also determined (546 OTUs identified). Several genera predominant in sow milk microbiota are commonly found in HM microbiota. They were also present in the SynComs, including *Staphylococcus*, *Streptococcus*, *Lactobacillus*, *Rothia* and *Corynebacterium* (Table *‘OTUs identified in sow milk’* accessible in the repository https://doi.org/10.57745/ATTHZ3). However, a few differences occurred between sow milk and HM microbiota (30, 31) or SynComs, such as the low abundance of *Bifidobacterium* (corresponding to less than 0.1 % of the total OTUs) or the presence of *Terrisporobacter* in sow milk.

The ileal, colonic and fecal microbiota of SM-fed piglets showed significant differences in both α-and β-diversity compared to the three formula-fed groups (Supplemental Figure S2), in agreement with differences in the relative abundance of numerous bacterial taxa (Supplemental Table S4). Interestingly, the SynCom-supplemented groups exhibited shifts in bacterial abundance compared to CTRL that aligned more closely with the SM group (Table 2 and Supplemental Table S4). This includes the higher abundance of Peptostreptococcaceae, Erysipelotrichaceae, *Clostridium sensu stricto 1*, *Turicibacter* in the ileum of HI, the higher abundance of *Alistipes* and lower abundance of *Butyricicoccus* in the colon and/or feces of AI. At PND8, the higher abundance of Prevotellaceae/*Prevotella* and lower abundance of Enterococcaceae/*Enterococcus* in the feces of HI and AI piglets compared to the CTRL group, was closer to levels observed in SM piglets.

To determine whether SynCom supplementation affected microbiota activity, SCFA concentrations were also analysed at PND8 (feces) and PND24 (feces and colonic digesta) (Table 3). The fecal concentration of acetate tended to be higher in HI compared to CTRL and AI and that of propionate was higher in HI compared to AI at PND8, whereas difference no longer occurred in feces at PND24 (Table 3). In the colon, propionate, valerate, butyrate, isovalerate, isobutyrate and total SCFA concentrations were higher in SM-fed than in formula-fed groups at PND24 (Table 3). Overall, adding HM-derived SynComs to the formulas specifically altered the microbiota composition. These changes reduced, but did not eliminate, the differences observed between CTRL and SM piglets. The effect of SynComs on fermentative activity was mainly observed in the first days of supplementation.

**Table 3.**
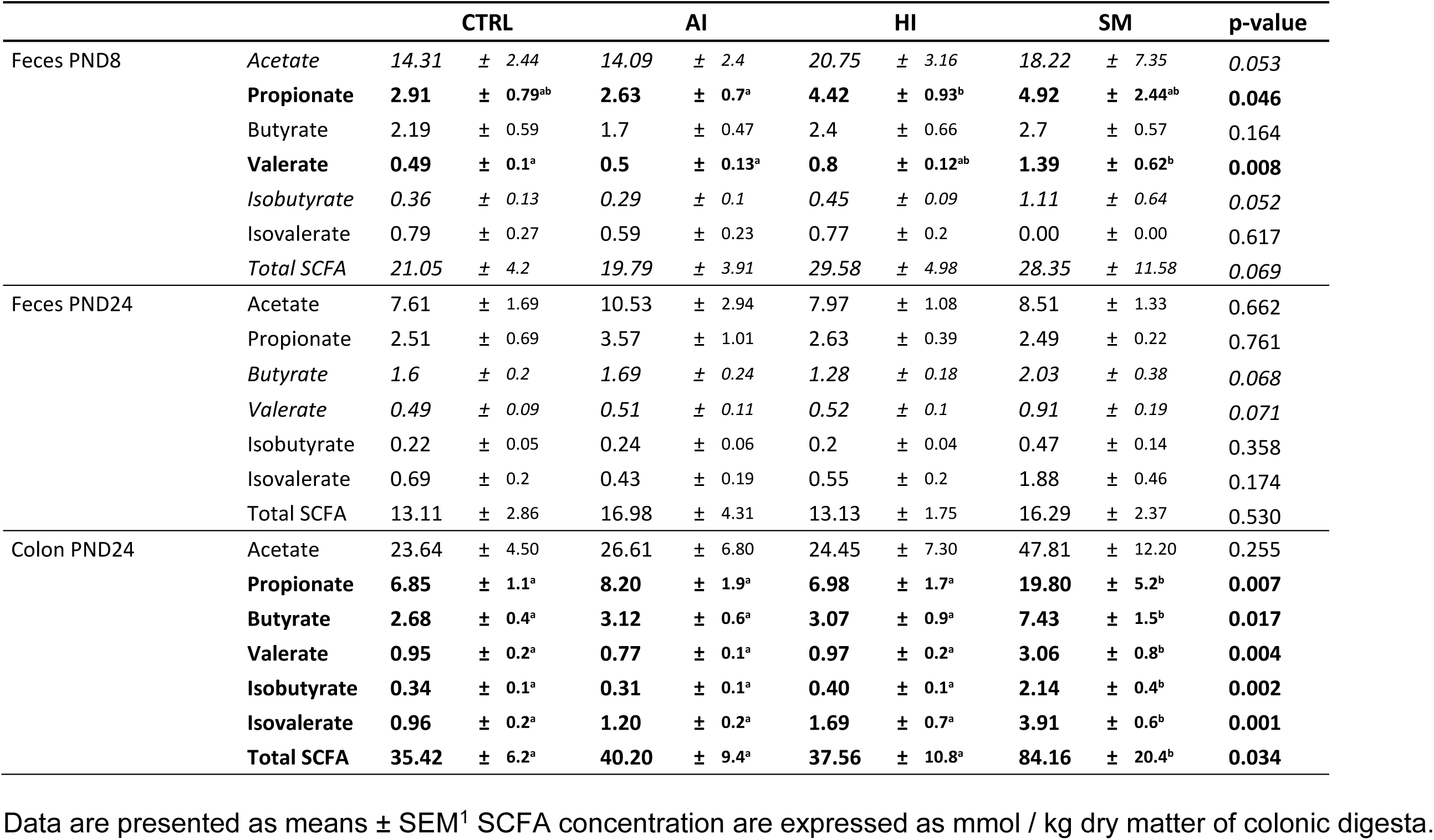
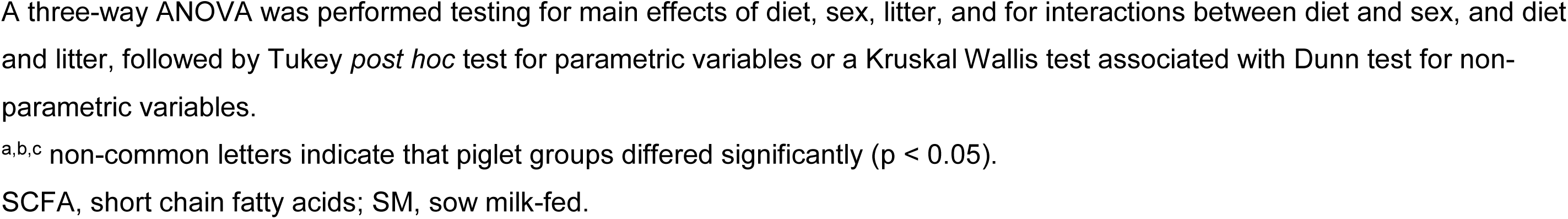
SCFA concentrations (mmol/kg dry matter) in the feces at PND8 and in the colonic digesta and feces at PND24 of formula-fed (CTRL, AI and HI) and sow milk-fed (SM) piglets.

|  |  | CTRL |  | AI |  | HI |  | SM |  | p-value |
| --- | --- | --- | --- | --- | --- | --- | --- | --- | --- | --- |
| Feces PND8 | Acetate | 14.31 | ± 2.44 | 14.09 | ± 2.4 | 20.75 | ± 3.16 | 18.22 | ± 7.35 | 0.053 |
|  | **Propionate** | **2.91** | ± 0.79<sup>ab</sup> | **2.63** | ± 0.7<sup>a</sup> | **4.42** | ± 0.93<sup>b</sup> | **4.92** | ± 2.44<sup>ab</sup> | **0.046** |
|  | Butyrate | 2.19 | ± 0.59 | 1.7 | ± 0.47 | 2.4 | ± 0.66 | 2.7 | ± 0.57 | 0.164 |
|  | **Valerate** | **0.49** | ± 0.1<sup>a</sup> | **0.5** | ± 0.13<sup>a</sup> | **0.8** | ± 0.12<sup>ab</sup> | **1.39** | ± 0.62<sup>b</sup> | **0.008** |
|  | *Isobutyrate* | 0.36 | ± 0.13 | 0.29 | ± 0.1 | 0.45 | ± 0.09 | 1.11 | ± 0.64 | 0.052 |
|  | Isovalerate | 0.79 | ± 0.27 | 0.59 | ± 0.23 | 0.77 | ± 0.2 | 0.00 | ± 0.00 | 0.617 |
|  | *Total SCFA* | 21.05 | ± 4.2 | 19.79 | ± 3.91 | 29.58 | ± 4.98 | 28.35 | ± 11.58 | 0.069 |
| Feces PND24 | Acetate | 7.61 | ± 1.69 | 10.53 | ± 2.94 | 7.97 | ± 1.08 | 8.51 | ± 1.33 | 0.662 |
|  | Propionate | 2.51 | ± 0.69 | 3.57 | ± 1.01 | 2.63 | ± 0.39 | 2.49 | ± 0.22 | 0.761 |
|  | *Butyrate* | 1.6 | ± 0.2 | 1.69 | ± 0.24 | 1.28 | ± 0.18 | 2.03 | ± 0.38 | 0.068 |
|  | *Valerate* | 0.49 | ± 0.09 | 0.51 | ± 0.11 | 0.52 | ± 0.1 | 0.91 | ± 0.19 | 0.071 |
|  | *Isobutyrate* | 0.22 | ± 0.05 | 0.24 | ± 0.06 | 0.2 | ± 0.04 | 0.47 | ± 0.14 | 0.358 |
|  | Isovalerate | 0.69 | ± 0.2 | 0.43 | ± 0.19 | 0.55 | ± 0.2 | 1.88 | ± 0.46 | 0.174 |
|  | *Total SCFA* | 13.11 | ± 2.86 | 16.98 | ± 4.31 | 13.13 | ± 1.75 | 16.29 | ± 2.37 | 0.530 |
| Colon PND24 | Acetate | 23.64 | ± 4.50 | 26.61 | ± 6.80 | 24.45 | ± 7.30 | 47.81 | ± 12.20 | 0.255 |
|  | **Propionate** | **6.85** | ± 1.1<sup>a</sup> | **8.20** | ± 1.9<sup>a</sup> | **6.98** | ± 1.7<sup>a</sup> | **19.80** | ± 5.2<sup>b</sup> | **0.007** |
|  | **Butyrate** | **2.68** | ± 0.4<sup>a</sup> | **3.12** | ± 0.6<sup>a</sup> | **3.07** | ± 0.9<sup>a</sup> | **7.43** | ± 1.5<sup>b</sup> | **0.017** |
|  | **Valerate** | **0.95** | ± 0.2<sup>a</sup> | **0.77** | ± 0.1<sup>a</sup> | **0.97** | ± 0.2<sup>a</sup> | **3.06** | ± 0.8<sup>b</sup> | **0.004** |
|  | **Isobutyrate** | **0.34** | ± 0.1<sup>a</sup> | **0.31** | ± 0.1<sup>a</sup> | **0.40** | ± 0.1<sup>a</sup> | **2.14** | ± 0.4<sup>b</sup> | **0.002** |
|  | **Isovalerate** | **0.96** | ± 0.2<sup>a</sup> | **1.20** | ± 0.2<sup>a</sup> | **1.69** | ± 0.7<sup>a</sup> | **3.91** | ± 0.6<sup>b</sup> | **0.001** |
|  | **Total SCFA** | **35.42** | ± 6.2<sup>a</sup> | **40.20** | ± 9.4<sup>a</sup> | **37.56** | ± 10.8<sup>a</sup> | **84.16** | ± 20.4<sup>b</sup> | **0.034** |

Data are presented as means ± SEM<sup>1</sup> SCFA concentration are expressed as mmol / kg dry matter of colonic digesta.

### SynCom supplementation affected systemic and intestinal immune functions

MFA indicated that immunity was one of the main functions affected by SynCom supplementation that influenced both the systemic and intestinal immune functions (Figures 4 and 5, Supplemental Tables S5 and S6). At PND8, the fecal sIgA content in HI piglets was 6- and 11-fold higher than in CTRL and AI piglets, respectively (Figure 4). A similar profile of sIgA content was observed in the ileal digesta at PND24, although the differences were non-significant. Meanwhile, SM piglets exhibited fecal sIgA levels 12-fold higher than the CTRL group at PND8, but only 1.9-fold higher than the HI group. At PND24, there was a trend toward higher cytokine secretion in HI peripheral blood mononuclear cells (PBMC) compared to CTRL and AI PBMC (Figure 4). A similar pattern was observed in PP from HI and AI piglets compared to the CTRL group (Supplemental Table S5). The impact of SynComs on intestinal immune function was further demonstrated through changes in gene expression in both ileal and colonic tissues (Figure 5). AI and HI groups showed a slight upregulation of genes associated with pro-inflammatory responses (CCL2, TNFa, IL4, IFNg, IL6, IL1b, IL8, CCL20), anti-inflammatory pathways (SOCS3), antioxidant activity (SOD2), Treg pathways (FOXP3), and cellular signalling (TLR2 and TLR4) compared to the CTRL group.

**Figure 4.**
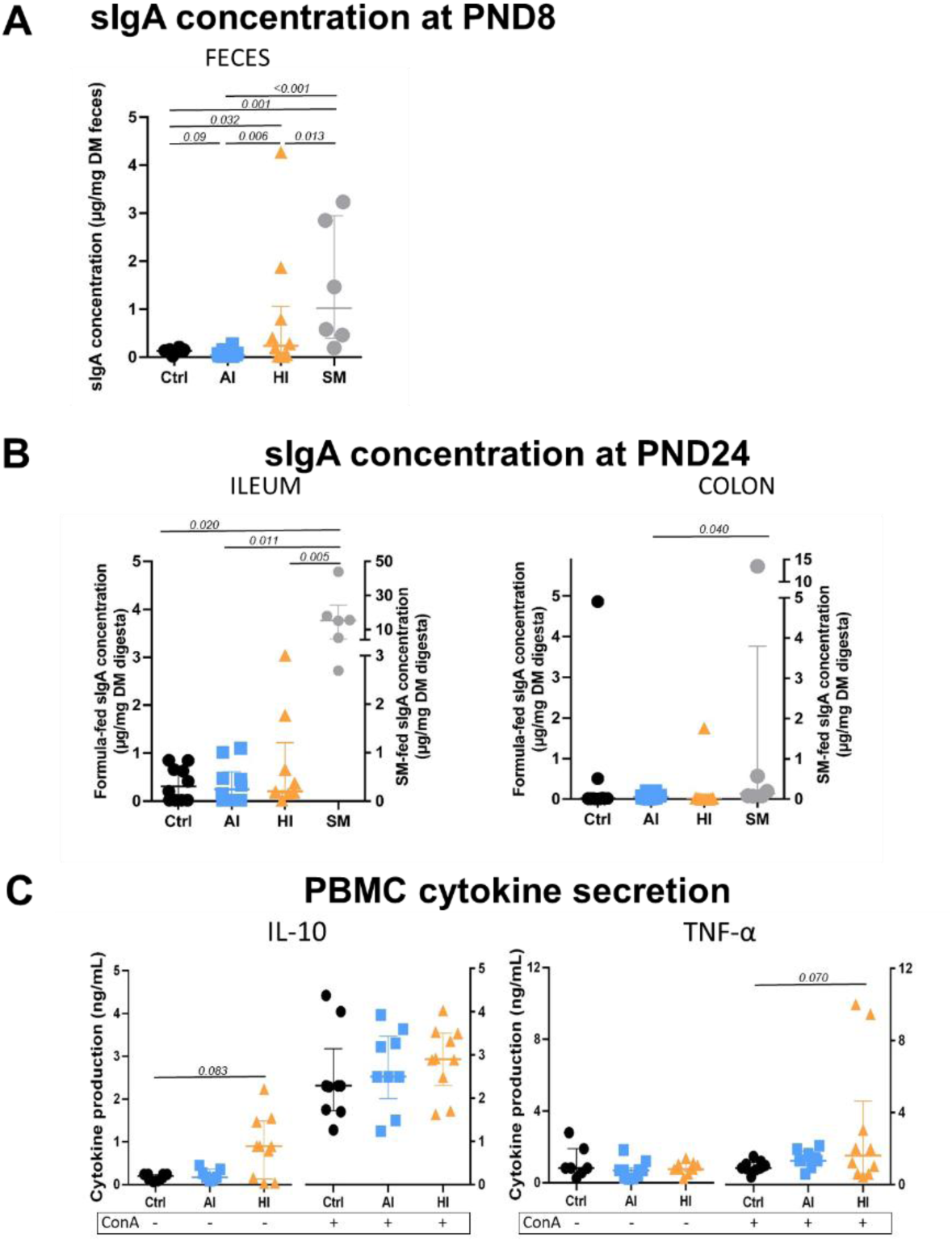
Secretory IgA (sIgA) concentration **(A)** in feces at PND8 and **(B)** in ileal and colonic digesta at PND24 of formula-fed (CTRL, AI, HI) and sow milk-fed (SM) piglets; **(C)** IL-10 and TNFα production in unstimulated (basal) and stimulated (Concanavalin A, ConA) conditions of formula-fed (CTRL, AI, HI) PBMC. Each individual is represented with one dot; Median with interquartile range. A three-way ANOVA was performed testing for main effects of diet, sex, litter, and for interactions between diet and sex, and diet and litter, followed by Tukey *post hoc* test for parametric variables or a Kruskal Wallis test associated with Dunn test for non-parametric variables. PBMC, Peripheral Blood Mononuclear Cells; PND, postnatal days

**Figure 5.**
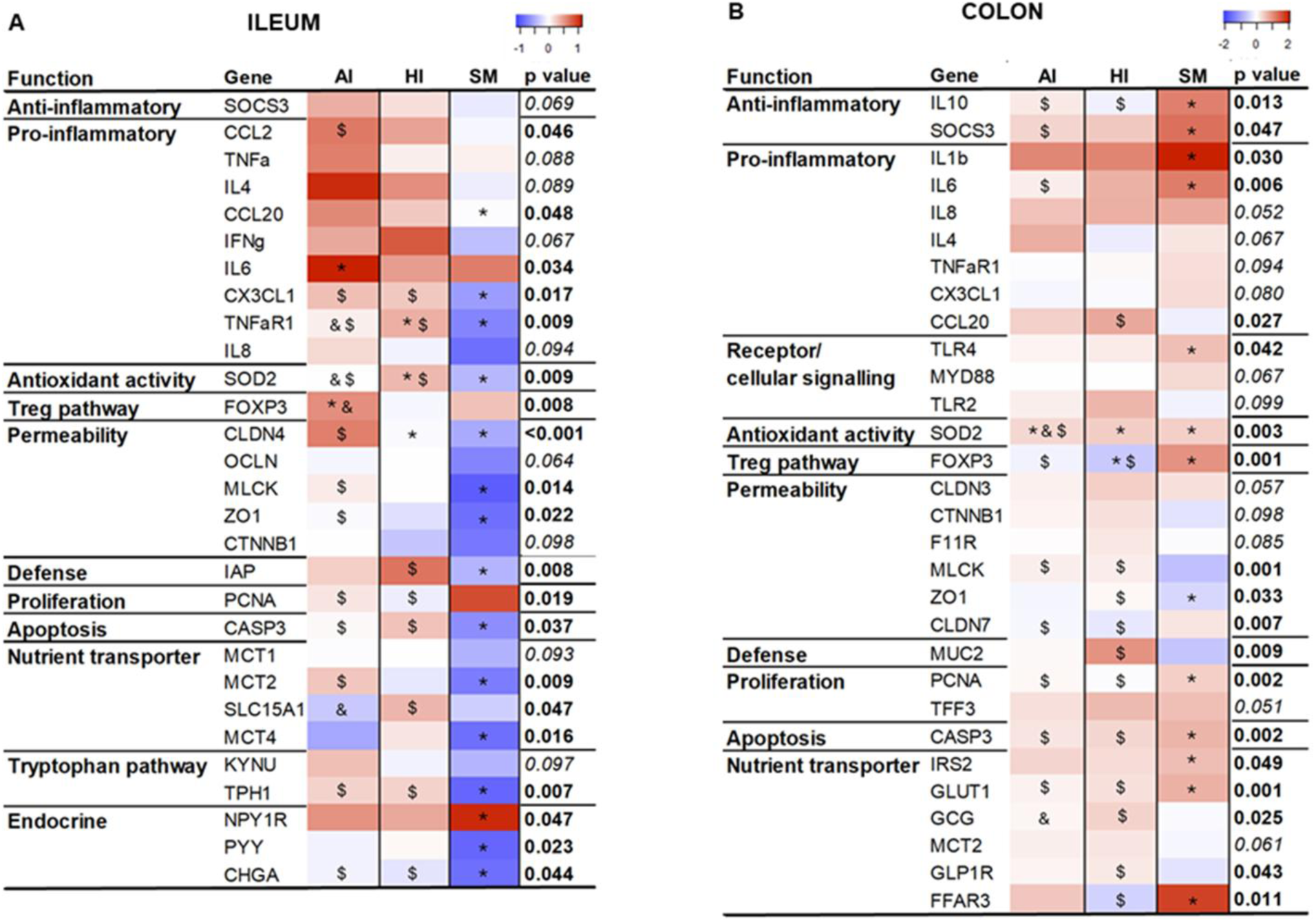
Heatmap of differentially expressed genes related to immune, epithelial barrier, nutrient transport, tryptophan metabolism and endocrine functions in ileum (A) and colon (B) of formula-fed AI and HI piglets and sow milk-fed (SM) piglets at PND24. The expressions of the target genes relative to the CTRL values were determined using the 2-ΔΔCt method for group comparisons. Gene expression data shown in the figure are log2 transformed data. A three-way ANOVA was performed testing for main effects of diet, sex, litter, and for interactions between diet and sex, and diet and litter, followed by Tukey *post hoc* test. Differences between the CTRL, AI, HI and SM groups were considered as statistically significant for p < 0.05 and a trend for difference at 0.05 < p < 0.1. When significant differences occurred, * indicates difference from CTRL, & difference from HI, and $ difference from SM. The full name of genes is given in the Supplemental Table S7. PND, postnatal days

Interestingly, the SM group also showed a mild stimulation of immune-related genes in the colon compared to the CTRL group, whereas the ileum exhibited an overall decrease in gene expression (Figure 5). Overall, adding SynCom HI -and SynCom AI to a lesser extent- to the formula modulated systemic and intestinal immune functions, and sIgA level specifically.

### Intestinal morphology and barrier functions differed between groups at PND24

To assess the effect of SynCom supplementation on the epithelial barrier, the intestinal morphology, the expression of genes related to cell renewal and barrier function, and *ex vivo* para- and transcellular permeability were evaluated in the ileum and colon at PND24 (Figures 5 and 6, Supplemental Table S6). The main differences were observed in villous height, goblet cell density, and genes involved in proliferation, apoptosis and mucin synthesis, particularly when comparing formula-fed piglets to SM piglets. Only minor differences were detected among the formula-fed groups (Figures 5 and 6).

**Figure 6.**
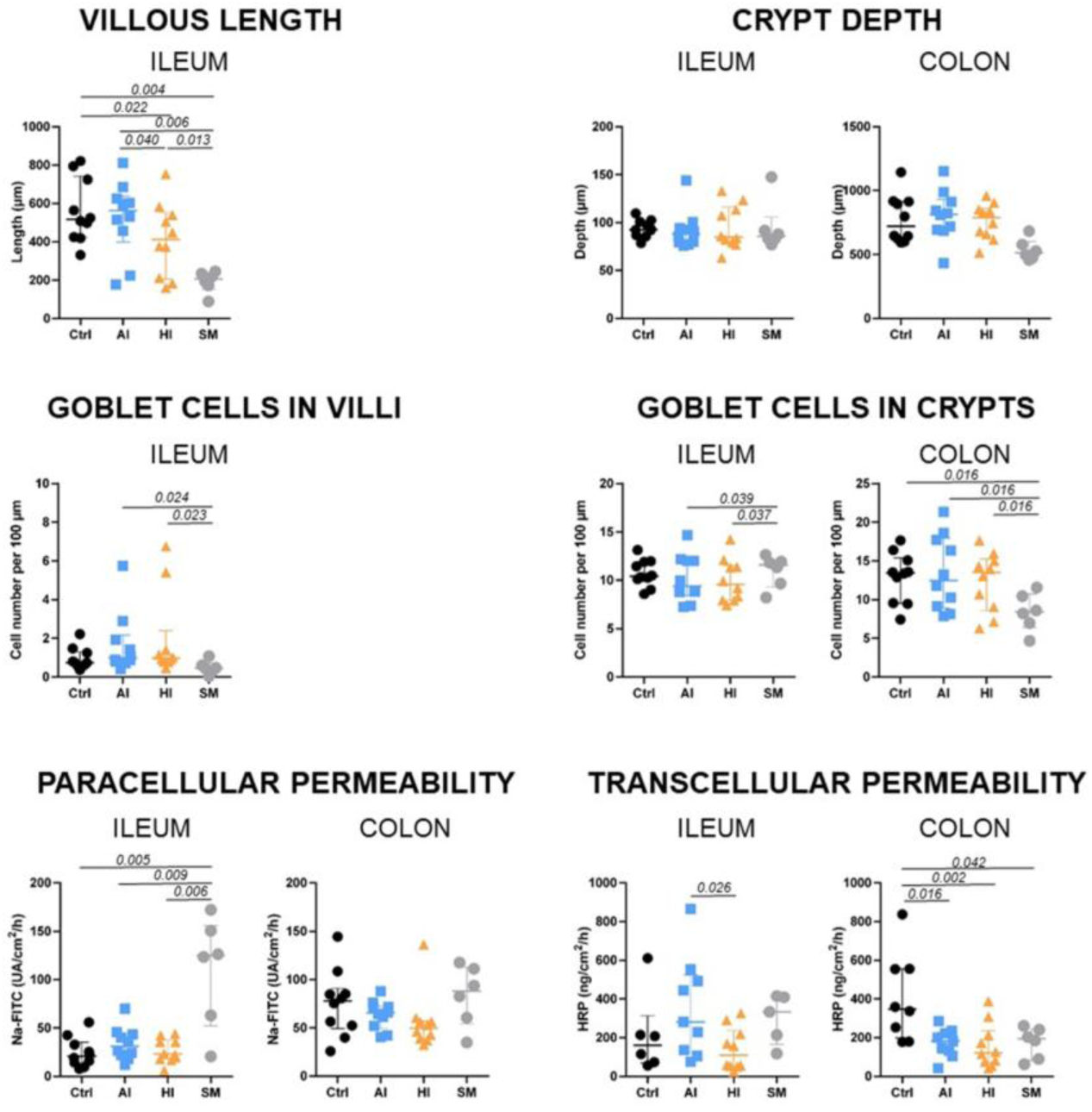
Effects of diets on intestinal barrier. Villous length in ileum and crypt depth in ileum and colon; goblet cell density in ileal villi and ileal and colonic crypts; *ex vivo* (Ussing chamber) epithelial barrier paracellular and transcellular permeability in ileum and colon, in formula-fed (CTRL, AI, HI) and sow milk-fed (SM) piglets at PND24. Each individual is represented with one dot; Median with interquartile range. A three-way ANOVA was performed testing for main effects of diet, sex, litter, and for interactions between diet and sex, and diet and litter, followed by Tukey *post hoc* test for parametric variables or a Kruskal Wallis test associated with Dunn test for non-parametric variables.PND, postnatal days

Regarding permeability, ileal paracellular permeability was lower in formula-fed piglets than in SM-fed piglets (Figure 6). This was accompanied by higher expression of ileal genes encoding tight junction proteins (CLDN4, MLCK and ZO1), as well as a trend toward increased expression of CTNNB1 and OCLN in the CTRL and AI groups compared to the SM group. The HI group showed intermediate expression levels (Figure 5A).

In the colon, there were no differences in *ex vivo* paracellular permeability among the four groups (Figure 6), despite some variations in tight junction protein expression (Figure 5B). Besides, transcellular permeability was higher in the ileum of the AI group compared to the HI group, while in the colon, both AI and HI groups exhibited lower transcellular permeability than the CTRL group, reaching levels similar to those in the SM group (Figure 6).

Overall, these results highlight the impact of SynCom supplementation on intestinal epithelium morphology and transcellular permeability.

### Intestinal nutrient transport, tryptophan metabolism and endocrine functions were moderately affected by the SynCom supplementation of formulas at PND24

A few number of genes related to nutrient transport, tryptophan metabolism and endocrine functions showed differential expression among formula-fed piglets (Figure 5 and Supplemental Table S6). Specifically, the ileal expression of SLC15A1 (peptide transporter) and the colonic expression of GCG (involved in endocrine function) were higher in the HI group compared to the AI group (Figure 5). Additionally, the expression of several other genes -encoding proteins involved in nutrient transport, tryptophan metabolism and endocrine function- varied between formula-fed and SM-fed piglets in both in the ileum and colon (Figure 5).

#### Correlation analyses highlight the influence of HM bacteria on the composition of the intestinal microbiota and intestinal functions

To explore individual associations between variables, correlation analyses were conducted between SynCom OTUs and microbiota or physiological variables (Figure 7), as well as between microbial genera and physiological variables (Supplemental Figure S3), specifically in the ileum and colon of formula-fed piglets.

**Figure 7.**
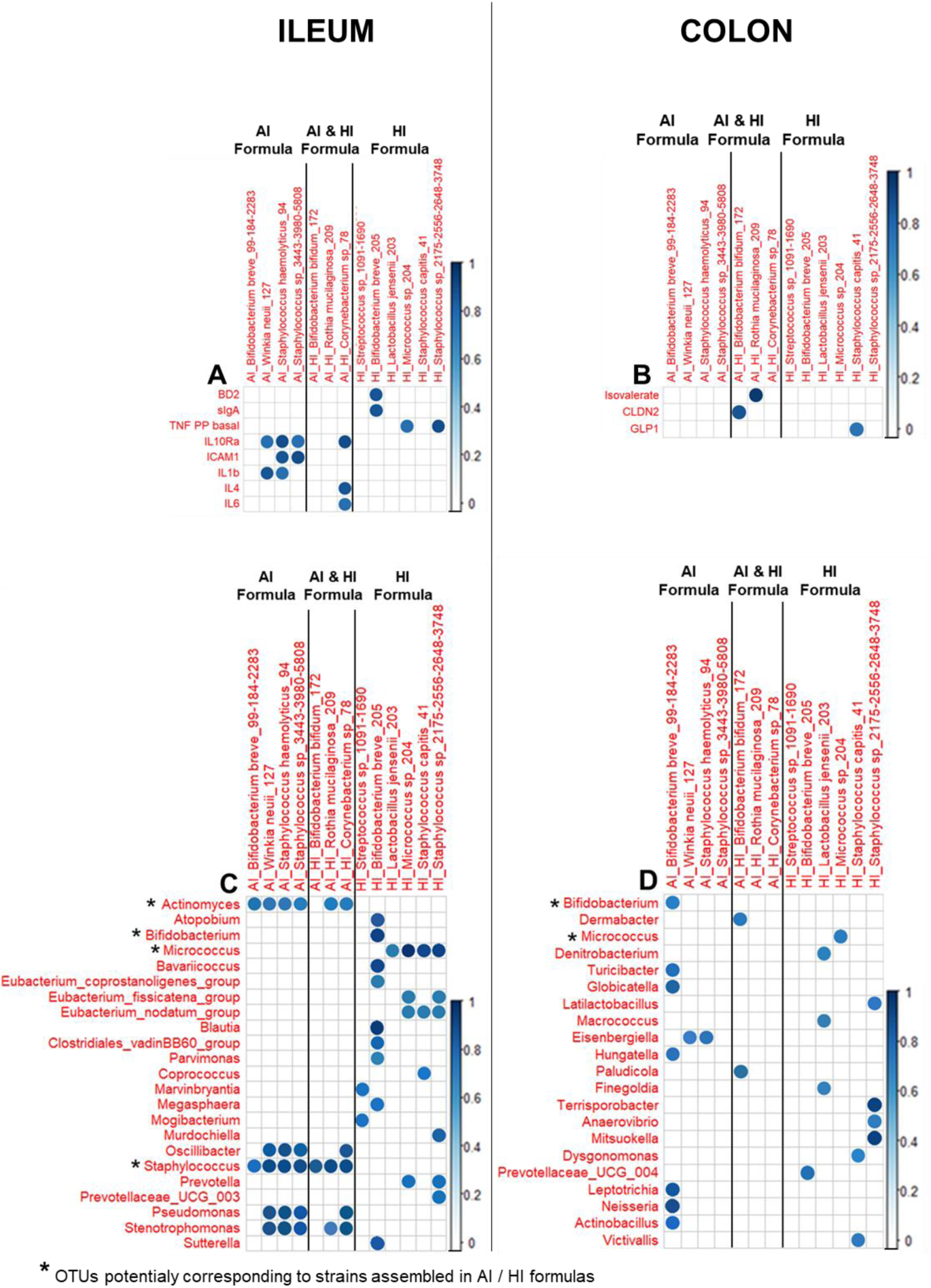
Heatmap of significant correlations with Benjamini–Hochberg correction showing the relationship between the relative abundance of the OTUs potentially corresponding to AI and/or HI SynCom strains, and **(A and B)** physiological variables and **(C and D)** microbial genus abundances in the **(A and C)** ileum and **(B and D)** colon of formula-fed piglets. Only associations with a correlation coefficient |r|> 0.7 and a significant p value < 0.05 are shown, with the strength of those correlations denoted by the intensity of the circle’s blue colour. Only positive correlations were significant. TNFα Basal PEYER, TNFα secretion by Peyer Patch cells in unstimulated condition ; IL10 LPS PEYER, IL-10 secretion by Peyer Patch cells in LPS-stimulated condition ; isovalerate, isovalerate concentration; all the other variables in capital letters correspond to the expression of genes listed in Supplemented Table S7.

In the ileum, the abundance of seven out of thirteen detected SynCom OTUs in the piglet microbiota showed strong correlations with eight immune-related variables. Notably, a specific OTU of the HI group corresponding to *Bifidobacterium breve*- was correlated to ileal sIgA level and to an ileal barrier variable (BD2) (Figure 7A). In contrast, only three SynCom OTUs correlated to a physiological variable in the colon (Figure 7B).

Additionally, all SynCom OTUs correlated to various genera in the ileum and eight SynCom OTUs correlated to various genera in the colon (Figure 7C and 7D).

Furthermore, multiple genera within the ileal and colonic microbiota exhibited significant positive or negative correlations (p < 0.05 and |r| > 0.7) with 37.1 % of total physiological variables in the ileum and 59.7 % in the colon (Supplemental Figure S3). These results underscore the influence of HM bacteria-driven gut microbiota composition on diverse gut functions.

## Discussion

Our study aimed to assess the impact of two SynComs on gut physiology and microbiota in piglets used as a model for human infants. The SynComs shared a similar taxonomic composition, reflecting the diversity of HM bacteria, but exhibited different immunomodulatory properties (9). From PND2 to PND24, piglets were fed with either a control formula or a formula supplemented with SynCom AI or HI at a constant concentration of 5.5 x 10^5^ CFU/mL (∼5 x 10^4^ CFU/mL per strain), comparable to the HM bacterial load in breastfed infants. Our findings indicate that most SynCom bacteria were present in the intestine of piglets fed the SynCom-supplemented formulas. The two SynComs exerted distinct effects on intestinal immune and barrier functions and microbiota composition, at both PND8 and PND24. SynCom HI showed a global impact on intestinal immune markers (including higher sIgA levels and PBMC cytokine secretion) associated with changes in microbial composition, particularly within the Bacillota phylum. The immune effects of the SynCom AI were less pronounced compared to SynCom HI. Correlation analyses further revealed associations between specific SynCom bacteria and both microbiota composition and intestinal functions, primarily in the ileum. Additional correlations between intestinal functions and genera not provided by the SynComs were mainly observed in the colon. This suggests that SynCom supplementation in formulas also induced physiological changes by altering the overall microbiota composition.

The HM bacteria assembled in the SynComs proved safe, causing no intestinal disorders or growth differences in piglets across the three formula-fed groups. OTUs corresponding to most added SynCom bacteria (“SynCom OTUs”) were detected in the ileum and, to a lesser extent, in the colon of SynCom-supplemented piglets, at a relative abundance of 0.0002 to 0.86 %. This aligned with evidence of vertical transfer of HM bacteria to the infant gut and the presence of probiotic bacteria in feces during supplementation (10, 32–35). We cannot exclude that these SynCom OTUs, when found in the intestinal microbiota, corresponded in part to bacteria others than SynCom bacteria, as metabarcoding does not allow to specifically identify a single strain. PCR with strain-specific primers or shotgun sequencing could undoubtedly confirm their identification. Nevertheless, their increased relative abundance in the ileum and colon of SynCom-supplemented piglets -compared to CTRL- strongly suggests that at least some of these OTUs originated from the added SynCom bacteria. No SynCom OTUs were found in the formula or in the intestine for *Cutibacterium*, *Streptococcus infantis,* and V*eillonella* strains. This may be due to their low abundance, bias in bacteria lysis or limitations of the primers used, which were suboptimal for amplifying the 16S rRNA gene of *Cutibacterium* and *Veillonella* (36). While the metabarcoding approach confirmed the presence of most SynCom OTUs in the intestinal microbiota of SynCom-supplemented piglets, it did not clarify whether these bacteria could survive or persist in the mucosal environment-especially since piglets received daily SynCom supplementation throughout the experiment. Of note, previous *in vitro* studies, however, demonstrated strain-dependent survival of five SynCom bacteria (namely *Bifidobacterium breve* CIRM-BIA2845, *Streptococcus salivarius* CIRM-BIA 2846, *Cutibacterium acnes* CIRM -BIA 2849, *Staphylococcus epidermidis* CIRM-BP 1633 and *Lactobacillus gasseri* CIRM-BIA 2841) in a model of infant gastrointestinal digestion (37).

The observed impact of SynCom OTUs on intestinal immune function -and, to a limited extent, on barrier function- underscores the modulatory influence of HM-derived bacteria on intestinal physiology. This effect is notable despite the low concentration of SynCom OTUs in the formulas and their limited relative abundance in the intestine. Since microbiota is less abundant in the ileum than in the colon (38), the relative abundance of the SynCom OTUs was higher in the ileum, where much of the gut-associated immune system resides. In this environment, where bacterial competition is reduced, even a low bacterial load may be sufficient to interact with the mucosa and its immune system. Previous studies have demonstrated similar immune-modulating effects at low doses. For instance, supplementing infant formula with either a low (10^4^ CFU/g of powder) or regular (10^7^ CFU/g of powder) dose of *Bifidobacterium lactis* produced comparable immune benefits in infants aged 0-6 months, despite a lower fecal detection of *Bifidobacterium lactis* in the low-dose group (39). Additionally, early-life variations in the abundance of subdominant bacterial genera of the gut mirobiota (such as *Lachnospira*, *Rothia*, *Veillonella* and *Faecalibacterium*) have been shown to reduce lung inflammation in an asthmatic mouse model (84), further supporting the potential of low-abundance bacteria to exert significant physiological effects. More generally, the ability of some species to influence the microbiota despite their low abundance is highlighted in the concept of keystone species that possess specific enzymatic activities, enabling, for instance, crossfeeding with others, sometimes more dominant, species, as illustrated for *Bifidobacterium* (85,86).

In line with the observed impact of SynCom OTUs on intestinal immune functions, seven of the thirteen detected SynCom OTUs in the piglet microbiota were correlated with ileal immune markers. These findings align with the immunomodulatory profiles of SynCom bacteria previously characterized *in vitro* (9). The correlations involved OTUs shared by or specific to SynComs AI and HI, including *Bifidobacterium breve, Corynebacterium sp*., *Staphylococcus haemolyticus, Winkia neuii* and *Micrococcus luteus*. Our results support the well-documented immunomodulatory role of *Bifidobacterium* strains as probiotics (40–44) and contribute to the limited existing knowledge about the other HM genera, highlighting their potential value for probiotic applications when assembled in SynComs.

Beyond the presence of SynCom OTUs in the intestine, SynCom supplementation also induced changes in the microbiota. SynCom HI supplementation, in particular, induced a higher ileal abundance of several families and genera mostly belonging to Bacillota, including *Clostridium sensu stricto 1*, *Eubacterium* and *Turicibacter*. This also included Peptostreptococcaceae, one of the most abundant families in the ileum (mean relative abundance >1% and up to >10%), with the AI group showing intermediate levels between HI and CTRL groups. In the colon, fewer changes were observed, mostly within Bacillota as well. Additionally, the AI group showed an increase of *Alistipes* compared to other groups.

Early immune boost reported in breastfed infants and HM-fed piglets was associated with the composition of the microbiota (5, 45–48). The role of Bacillota in immune ontogeny has been demonstrated in rodents, where it is essential for regulating susceptibility to immunopathologies later in life (49). *Turicibacter*, a known indicator of immune status, was positively correlated with the TNFα response of ileal PP cells in this study. Its abundance was decreased in immunodeficient mice (50, 51) and negatively associated with immune-related adverse events (52). *Alistipes*, a human gut commensal involved in protein fermentation, has been linked to immune modulation, though its effects remain debated (53, 54). Some species, such as *Alistipes onderdonkii*, have shown potential as anti-inflammatory probiotics (55). Correlation analyses further revealed that specific SynCom OTUs, such as *Bifidobacterium breve* (OTU 205), were directly associated with ileal sIgA levels, a critical marker of immune development in infants. This SynCom OTU also correlated with other genera, including *Bavaricoccus* and *Parvimonas* (Bacillota) and *Sutterella* (Pseudomonadota), some of them being likewise linked to sIgA levels. sIgA plays an important role in the intestinal defense by preventing pathogen adhesion and neutralizing toxins. Notably, fecal sIgA levels were higher in the HI group at PND8, suggesting a beneficial effect of Syncom HI-induced microbiota changes on immune markers. These findings align with previous studies showing that probiotic supplementation can restore sIgA levels in formula-fed infants, which are typically lower compared to breastfed infants (56–58).

Despite modulation by SynCom-supplementation, the microbiota still differed between the three groups of piglets fed formula and SM. Specifically, microbial α-diversity was lower in the SM group compared to the formula groups. This aligns with the lower α-diversity observed in the feces of infants fed HM compared to formula at 1-2 months of age (59–65) or in the ileum, colon and/or rectum of 3-week-old piglets fed HM compared to formula (5, 66, 67). However, no difference in intestinal α-diversity was found in weaned piglets fed sow milk or milk replacer (68). β-diversity also differed in piglets fed SM or formula, consistent with previous studies comparing formula with SM or HM (5, 66). These differences may be linked to the contrasting environments of piglets, including maternal presence, as well as the specific role of maternal milk components like oligosaccharides. In line with the differences in β-diversity, several taxa showed differential abundances between formula-fed and SM-fed piglets. Yet, some of the changes induced by Syncom HI -and, to a lesser extent, AI-supplementation, compared to the CTRL group, brought the SynCom-supplemented groups closer to the SM group. This includes the higher abundance of Bacillota members in the ileum of the HI group and of *Alistipes* in the colon of the AI group, as well as the higher abundance of Prevotellaceae/*Prevotella* and lower abundance of Enterococcaceae*/Enterococcus* at PND8 in the feces of HI and AI compared to the CTRL piglets. Notably, a rapid decrease in Enterococcaceae after birth has been reported during the maturation of piglet colonic microbiota, while Prevotellaceae/*Prevotella* was more abundant in HM-fed piglets compared to formula-fed ones (5, 17, 66). Furthermore, several taxa exhibited differential abundance in the HI and AI groups compared to the CTRL group at PND8, even more so than at PND24. This was associated with changes in fecal metabolite concentrations in the HI group, particularly higher acetate and propionate levels compared to the CTRL and AI groups at PND8, resembling those observed in the SM group. SCFAs may play a role in immune maturation, both locally in the intestine and systemically (69). Overall, these results suggest that SynCom HI, and to a lesser extent AI, may drive changes in microbiota composition and activity toward that of SM-fed piglets.

The correlations between certain genera -whose abundance was either altered or unchanged by SynCom supplementation- and ileal and colonic barrier variables (such as genes encoding tight junction proteins and mucins) highlight the relationship between microbiota and intestinal barrier functions, as previously observed (5). However, no differences in *ex vivo* paracellular permeability were found between the formula-fed groups in the ileum and colon. In line with this, the expression of genes encoding tight junction proteins showed little variations among formula-fed piglets, but was reduced in SM piglets, with HI piglets sometimes exhibiting intermediate levels. Notably, the higher paracellular permeability observed in the SM ileum aligns with findings from some studies in breastfed infants (70) and HM-fed piglets (5), which reported higher permeability compared to formula-fed piglets. Yet, these results remain controversial, as other studies have found no change -or even a reduction- in total gut permeability in breastfed infants compared to formula-fed infants (1, 71–73), or reduced expression of genes coding for tight junction proteins in HM-fed piglets (74). Additionally, changes in the expression of genes involved in other functions – such as nutrient transport, endocrine function and tryptophan metabolism- were significantly correlated with numerous genera in the ileum and colon microbiota. This further supports the idea that SynComs AI and HI influence intestinal functions by modulating the composition of the intestinal microbiota.

In conclusion, HM-derived bacteria can reach intestinal compartments and influence the intestinal epithelium, its immune function, and the microbiota. Consistent with our *in vitro* study using a quadricellular model mimicking the intestinal epithelium, we confirmed that SynComs -despite having similar taxonomic composition but contrasting immunomodulatory properties- can differentially modulate the developmental profile of intestinal immune functions. The immunostimulatory profile of SynCom HI induced more pronounced changes in intestinal immune properties than the anti-inflammatory profile of SynCom AI, as evidenced by effects on sIgA levels and cytokine secretion by PBMC. A major limitation of this study was that the SynComs, composed of only 11 strains, did not fully capture the complexity of the HM microbiota. Additionally, the strains were assembled in equal proportions, whereas dominant taxa are typically observed in the HM microbiota (4, 6). Nevertheless, this study demonstrates that the continuous dietary supply of HM-like SynComs at physiological doses -5.5 x 10^5^ CFU/mL, which is within the upper mid-range of HM bacterial load-was sufficient to induce physiological effects in a SynCom-dependent manner. Further investigation is needed to fully characterize underlying mechanisms and identify the keystone bacteria. Moreover, the apparently beneficial effects observed with SynCom OTUs in healthy piglets should be evaluated in unhealthy contexts where the immune system is challenged, such as allergic conditions, which are common in infants.

## Materials & Methods

### Animal study

The study was designed and conducted at INRAE experimental facilities (Pig Physiology and Phenotyping Experimental Facility, Saint-Gilles, France, doi 10.15454/1.5573932732039927E12; Agreement No. D35-275-32) in accordance with the current ethical standards of the European Community (Directive 2010/63/EU), and the French legislation on animal experimentation and ethics. The ethics committees of CREEA (Rennes Committee of Ethics in Animal Experimentation) and of France’s Ministry of Higher Education and Research approved (authorization #35727-2022030411197283) the entire procedure described in this protocol. Animal welfare was ensured by daily observation throughout the experiment. The piglets did not receive any medication or antibiotic treatment.

Thirty female and male Yucatan piglets were separated from their dam at postnatal day (PND) 2 and housed in individual stainless-steel metabolic cages. Room temperature was maintained at 30°C for the first week and then lowered at 28°C until the end of the experiment. Piglets were fed one of the three experimental formulas with an automatic milk feeder, as previously described (26). To account for litter-to-litter variation, three piglets with a BW close to the mean birth weight of the litter were selected from each litter and assigned to one of the three formulas. Mean birth weight of selected piglets was 0.849 ± 0.023 kg. Allocation to formulas was balanced between groups for birth weight, BW at PND2 and sex. Formulas were daily rehydrated to 20 % dry matter extract in water prior to distribution. Formula supply was divided into ten meals that were automatically distributed throughout the day. BW was measured twice a week and feeding amounts were adjusted accordingly. The daily net energy offered was 1450 kJ/kg metabolic BW. Formula intake was automatically recorded for each meal. In addition, six piglets selected from the same litters as the formula-fed piglets were allowed to suckle the sow naturally (SM) to provide benchmarks for gut development and microbiota. Seventeen female and nineteen male were used (CTRL: n=4 female and 6 male; AI: n=5 female and 5 male; HI: n=4 female and 6 male; SM: n=4 female and 2 male). All piglets were euthanized at PND24 (± 2.6 days) and tissues were collected.

### Diets and synthetic bacterial communities

Formulas were manufactured at a semi-industrial pilot scale at Bionov (Rennes, France). The three formulas had the same nutritional content (Supplemental Table S1). They differed by the bacterial supplementation: two HM-derived SynComs with anti-inflammatory (AI) *vs.* high immunomodulatory (HI) properties (9) as described in Table 1, or no bacterial supplementation for the control (CTRL) formula.

The AI and HI SynComs were each composed of 11 strains in equal proportions. Three of the strains were common to both SynComs and a further eight strains were specific to each SynCom. Firstly, the two SynComs were designed to mimic HM microbiota with representatives of genera frequently found in HM and with at least, one representative of the 4 most widespread genera of the cultivable HM microbiota: *Staphylococcus*, *Cutibacterium*, *Streptococcus* and *Corynebacterium* (6). Secondly, to consider the wide variability in the immunomodulatory capacities of HM bacteria, the 2 communities were assembled to have theoretically contrasting immunomodulatory effects, depending on their individual properties, as previously described (9). The SynCom AI was designed to display mainly anti-inflammatory properties and the SynCom HI to display both anti- and pro-inflammatory properties. Each bacterial strain was cultured separately on its optimal growth medium for 1 to 5 days as previously described (9) and its concentration was estimated with an OD measurement. Bacterial cells were harvested by centrifugation (6000 g, 10 min, 4°C), washed with 0.9 % (wt /vol) saline solution, pooled and resuspended in the formula. Daily doses equivalent to a 500-fold concentrated bacterial suspension (total bacterial concentration of 2.75 x 10^8^ Colony Forming Units (CFU)/mL) were prepared and stored at -80°C until used. After thawing in a water bath at 37°C for a few minutes, AI and HI bacterial doses were added in the AI and HI diets, respectively, at a total bacterial concentration of 5.5 x 10^5^ CFU/mL of formula (corresponding to a concentration of 5 x10^4^ CFU/mL of formula for each strain).

### Sample collection

#### Diet

Fifteen mL milk samples were collected from sows nursing the SM piglets on day 20 postpartum. Injection of oxytocin (2 mL, Laboratoires Biové, Arques, France) was carried out to facilitate milking. Before sampling, the sow teats were thoroughly washed with water and soap and rinsed with sterile physiological water (0.9 % NaCl) before cleaning with ethanol. Samples of formulas CTRL, AI and HI were also randomly collected at the time of rehydration of formula powders during the third week of the experimental period. Sow milk and formula samples were stored at -80°C until microbiota analysis.

#### Piglets

During the experimental period, feces were collected at PND8 and stored at -80°C for microbiota, SCFA and sIgA analyses. On the last day of the experiment, piglets were euthanized 1 h after their last meal by electrical stunning immediately followed by exsanguination. Blood was collected in 2 BD Vacutainer CPT (BDBiosciences, NJ, USA) and stored at room temperature until PBMC extraction. Ileal digesta and tissue were collected from an 80-cm segment anterior to the ileocecal junction, and colonic digesta and tissue were collected from the first third part of the colon. Samples of 100 mg ileal and colonic digesta were immediately frozen in liquid nitrogen and stored at - 80°C for microbiota and sIgA analyses. Samples of feces were collected and stored at -80°C for microbiota analysis. In addition, 0.5 to 2 g of ileal and colonic digesta and feces were collected for SCFA analysis. A 10-cm segment of distal ileum (containing the ileal PP) was collected, rinsed with ice-cold Hank’s balanced saline solution supplemented with 50 mg/mL gentamicin, 200 UI/mL penicillin, 200 mg/mL streptomycin and 10 mM hepes for isolation of PP mononuclear cells. The remaining ileal segment and the proximal colon were rinsed with cold phosphate buffered saline (PBS). A 10-cm segment was kept in ice-cold Dulbecco’s Minimum Essential Medium (Gibco, Thermo Fisher, France) for immediate Ussing chamber analysis. About 100 mg of ileal (without PP) and colonic tissues were kept in a RNA later solution for 24 h at 4°C and stored at −20°C until RNA extraction and gene expression analysis. Finally, adjacent segments (10-cm) were fixed in 4 % paraformaldehyde for 48 h until further dehydration in ethanol and embedding in paraffin, for morphometry analysis and goblet cell counting.

### Microbiota analysis

Extraction of total bacterial DNA from ileal and colonic digesta and feces was performed as described in the instruction guide of the Quick-DNA Fecal/Soil Microbe Miniprep Kit (ZYMO Research, Irvine, USA). Sow milk and formula samples were pre-treated before total bacterial DNA extraction as described (75). The V3-V4 region of 16S rRNA gene was amplified using the following primers: CTTTCCCTACACGACGCTCTTCCGATCTACTCCTACGGGAGGCAGCAG (V3F) and GGAGTTCAGACGTGTGCTCTTCCGATCTTACCAGGGTATCTAATCC (V4R), Phusion® High-Fidelity DNA Polymerase (New England Biolabs, Évry-Courcouronnes, France) and dNTP (New England Biolabs) for 25 cycles (10 s to 98°C, 30 s at 62°C, and 30 s at 72°C). Agarose gel electrophoresis was performed to verify amplicon purity prior to sequencing using Illumina MiSeq technology, performed at the Genotoul GeT-PlaGe platform (Toulouse, France).

#### SCFA analysis

After collection, ileal and colonic digesta and feces were weighed, mixed with 1 mL 0.5 % orthophosphoric acid solution per g of digesta and centrifuged at 1700 g for 15 min at 4°C. Supernatants were then stored at -20°C until SCFA quantification. SCFAs were analyzed by High Performance Liquid Chromatography (HPLC, Ultimate 3000, Thermo Fisher Scientific, Courtaboeuf, France). Acid separation was performed using a Rezek ROA organic acid H+ column (300*7.8mm Phenomenex, California) with H2SO4 0.005M as mobile phase at a flow rate of 0.4ml/min at 60°C. Two detectors were used: a UV detector (Dionex UVD 170U) operating at 210nm and a refractometer (RI 2031 Plus Jasco). Quantification was performed with an external calibration using acetic acid (PanReac, Lyon, France), propanoic, 2-methylpropanoic, butanoic, 3-methylbutanoic and pentanoic acids (Merck, St. Quentin Fallavier, France) as standards (5).

### *Ex vivo* permeability measurement

Ileal and colonic permeability measurements were performed using Ussing chambers (Physiological Instruments, San Diego, CA, USA) as described (5). Permeability was determined using tracer molecules, fluorescein sodium salt (Na-FITC) for paracellular permeability and peroxidase from horseradish type VI (HRP) for transcellular permeability. The tracer molecules were added into the apical compartment and those transferred through the epithelium were analyzed in the serosal compartment. Concentration of Na-FITC in the samples collected after 120-min incubation from the serosal buffer was measured by fluorimetry (Varioskan™ LUX multimode microplate reader, ThermoFisher Scientific, Saint-Herblain, France) whereas concentration of HRP was determined using spectrophotometry (Varioskan™ LUX multimode microplate reader, ThermoFisher Scientific, France) after enzymatic reaction using o-Dianisidine dihydrochloride as substrate (Merck, Molsheim, France).

#### Histomorphometry analysis

Histomorphometric analysis was performed after alcian blue and periodic acid Schiff staining on 7 μm sections of formalin-fixed and paraffin-embedded ileal and colonic tissues. Sections were analysed using a light microscope (Nikon Eclipse E400, Nikon Instruments, France) with image processing software (NIS-Elements AR 3.0, Nikon Instruments), as described (76). Villous and crypt sizes and goblet cell density were measured in at least 15-20 crypt-villous units per piglet.

### Cell isolation and *in vitro* culture

PBMC extraction was performed within 2 h after collection following the manufacturer instructions of the BD Vacutainer CPT (BDBiosciences, NJ, USA). After centrifuging the BD Vacutainer CPT tube at 1500 g for 20 min, the cell ring was recovered. The PBMC cells obtained were resuspended with 10 mL of HBSS supplemented with 10 % Foetal Calf Serum (FCS) and centrifuged 10 min at 600 g and 4°C. The supernatant was discarded. The pellet containing PBMC was resuspended with 7 mL of Ammonium-Chloride-Potassium (ACK) lysis buffer, and with 7 mL of RPMI complemented with 10 % FCS and 1 % Penicillin/Streptomycin (complete RPMI). The suspension was centrifuged 5 min at 600 g and 4°C. The pellet was resuspended with 14 mL of complete RPMI and centrifuged 10 min at 600 g and 4°C. Finally, PBMC were suspended in complete RPMI to achieve cell concentration of 4.10^6^ cells/mL. Mononuclear cells from freshly-removed ileal PP were isolated as previously described (76). PP cells were suspended in complete RPMI to achieve cell concentration of 8.10^6^ cells/mL.

PBMC and PP mononuclear cells were cultured in RPMI complete medium for 48 h at 37°C in a 5% CO_2_ water-saturated atmosphere. Culture was performed in unstimulated conditions and stimulated conditions with ConA (5 µg/ml of Concanavalin A from Canavalia ensiformis Merck, Molsheim, France) or LPS (10 µg/ml of LPS-EB ultrapure from Escherichia coli 0111:B4 strain, InvivoGen, San Diego, USA). In addition, PP mononuclear cells were cultured at 37°C in a 5 % CO_2_ water-saturated atmosphere for 7 days to assess sIgA production. Supernatants of PBMC and PP cells were collected and following the addition of antiprotease cocktail 1X (SigmaFast, Merck Sigma, Saint Quentin Fallavier, France) stored at −20 °C until cytokine and sIgA analyses.

#### Cytokine and sIgA analyses

Concentrations of IL-10 and TNF-α in culture supernatants of PBMC and PP cells were measured by ELISA (R&D Systems, USA: DY693B and DY985, respectively).

For sIgA analysis, ileal and colonic digesta at PND8 and PND24 and fecal samples at PND8 were 10-fold diluted with PBS-EDTA (0.5 M) buffer supplemented with a protease inhibitor cocktail (250 µg/L; P2714-1BTL). After a vigorous shaking, suspensions were incubated 30 min at room temperature on a Rotary laboratory shaker and centrifuged at 18 000 g, 30 min and 4°C. Supernatants were stored at - 20°C until ELISA analysis. Concentration of sIgA in 7 days-cultured PP cell supernatants as well as in ileal, colonic and fecal supernatants was assessed as previously described (76).

#### Gene expression analysis

Total RNA extraction from ileal and colonic tissues was performed using the "NucleoSpin® RNA" kit (Macherey Nagel). Extracted RNA was quantified using a DS-11 spectrophotometer (DeNovix, Wilmington, DE, USA). RNA quality and RNA Integrity Number (RIN) were assayed using the Agilent RNA 6000 Nano kit in combination with an Agilent 2100 Bioanalyzer (Agilent Technologies France, Massy, France). Reverse transcription was then carried out on 1 μg of extracted RNAs using the High Capacity Complementary DNA Reverse Transcription Kit, as previously described (77).

Quantitative real-time PCR was performed on a 384-well plates real-time PCR machine using a PowerSYBR green PCR Master Mix kit (4368708, ThermoFisher Scientific, France) for detection (78). The housekeeping genes selected were YWHAZ and PGK1 for ileum, and HTPR1 and RPL4 for colon. Relative expressions of the target genes (Supplemental Table S7) were determined using the 2^-ΔΔCt^ method for group comparisons.

## Statistical analysis

Data are presented as means ± SEM (standard error of the mean). For microbiota data, raw sequences were analyzed using the bioinformatic pipeline FROGS (Find Rapidly OTU with Galaxy Solution) software (79). Data were treated with the following steps: pre-processing, clustering with swarm algorithms, remove of chimera, OTU filters, and finally taxonomic affiliation. The descriptive analysis of the structure (α- and β-diversity) of the microbiota was conducted with the Phyloseq function (EdgeR package, Bioconductor). The α-diversity index used were Chao1 representing the bacterial richness. Phylogenetic β-diversity was studied using the Bray-Curtis distances and group differences were evaluated with principal co-ordinate analysis (PCoA) and permutational multivariate analysis (PERMANOVA) of variance using distance matrices. Differences in phyla, families, genera and OTUs were assessed with pairwise comparisons on OTUs or after aggregation at the desired taxonomic rank (phyloseq tax_glom function) using DESeq2 as described in (5).

*Unidimensional analysis.* A statistical analysis of the variables (excluding microbiota data) was performed using R software (version 4.3.2). Body weight differences between groups (SM, CTRL, AI and HI) were assessed using repeated-measures ANOVA, testing for the effects of diet, time and sex. For the other variables, a three-way ANOVA was performed testing for main effects of diet, sex, litter, and for interactions between diet and sex, and diet and litter, followed by Tukey *post hoc* test for parametric variables. When sex or litter effects or their interactions with diet were non-significant (p > 0.05), these factors were removed from the model. The normal distribution and the homoscedasticity of the residuals of each linear model were tested using Shapiro-Wilk and Levene’s tests, respectively. When the raw data did not fulfil these model assumptions, a natural logarithmic transformation of the data was performed prior to running the linear model. If the assumptions were still not satisfied, data were tested with a non-parametric Kruskal Wallis test associated with Dunn test. Differences were considered as statistically significant for p < 0.05 and a trend for difference at 0.05 < p < 0.1.

*Multidimensional analysis.* Partial Least Squares Discriminant Analyses (PLS-DA; mixOmics package (80)) were performed on all the variables measured in the ileum (n=201) and colon (n=250) to calculate their Variable Importance in Projection (VIP) score (81) on the first and the second components of the PLS-DA that discriminated the three formula groups. The variables with a VIP score greater than 1 on components 1 or 2 were selected, corresponding to 112 variables for the ileum and 141 variables for the colon. Variables were then divided into eleven groups as follows: Barrier, Immunity, Microbiota activity, Nutrient transporter, Endocrine and Tryptophan metabolism, plus five microbiota phyla: Actinomycetota, Bacillota, Bacteroidota, Pseudomonadota, and ’Other Phyla’ (that gathered the Campylobacterota, Deferribacterota, Desulfobacterota, Euryarchaeota, Fusobacteriota, Spirochaetota, Synergistota, Verrucomicrobiota and Patescibacteria phyla), for the Multifactorial analysis (MFA; FACTOMINE R package (82, 83)).

*Data correlation.* Pearson correlation coefficients with Benjamini–Hochberg correction were determined between the SynCom bacteria, and microbiota and physiological variables, and between microbial genera and physiological variables, in the ileum and colon of formula-fed piglets. Relevant significant correlations (|r| ≥ 0.7, p < 0.05) are presented in the Figure 7 and the Supplemental Figure S3.

## Supporting information

Supplemental Figure S1.jpg

Supplemental Figure S2.jpg

Supplemental Figure S3.jpg

## Acknowledgement

We warmly thank all volunteer mothers for their breastmilk samples and all the paediatric nurses of the maternity department of Rennes University Hospital, for their involvement in the project. The authors acknowledge the staff of Rennes Pig Physiology and Phenotyping Experimental Facility (UE1421 UE3P, Saint-Gilles, France, doi 10.15454/1.5573932732039927E12; Agreement No. D35-275-32) for their participation in the realization of this experiment and their expert assistance in animal care and feeding. We thank the Genotoul GeT-PlaGe platform from Toulouse (France) for 16S rRNA metabarcoding.

This work was funded by the Région Bretagne (grant no. 19008213) and Région Pays de la Loire (grant no. 2019-013227) (France), and by Bba Milk Valley association through the PROLIFIC project.

## Data Availability Statement

All data generated or analysed during this study are included in this published article and its supplementary information files. The datasets presented in this study can be found in online repositories (four Tables named ‘Relative abundance of the OTUs that did not correspond to AI and HI bacteria in formulas’, ‘Individual ileal and colonic data used for PLS-DA and MFA’, ‘OTUs identified in sow milk’, ‘Cytokine production by Peyer Patch and PBMC cells in formula-fed piglets’) https://doi.org/10.57745/ATTHZ3.

In addition, the raw sequences of microbiota analysis are available on NCBI repositories (PRJNA1197167).

## Contributions

ILHL, SE, YLL and SB conceived the project that led to the submission of this work. ILHL and SE supervised research; CLB, SE and ILHL designed the experiment; CLB, GR, AC, PD, SG, RJ, VR, LR, MBM, SE and ILHL acquired data; CLB, SE and ILHL played an important role in interpreting the results; AB and LR realised the human milk collection; CLB, SE and ILHL drafted the manuscript and had primary responsibility for final content. All authors revised and approved the final manuscript.

---

**Supplemental Figure S1.** Partial Least Squares Regression-Discriminant Analysis (PLS-DA) of **(A)** 201 microbial and physiological variables analyzed in the **ileum** and **(B)** 250 microbial and physiological variables analyzed in the **colon** of formula-fed (CRTL, AI and HI) piglets at PND24. The 30 most discriminating variables for Component 1 and Component 2 are listed on the figures; the complete list of discriminant variables is given in ‘the online dataverse https://doi.org/10.57745/ATTHZ3 - Table “Individual ileal and colonic data used for PLS-DA and MFA”. PND, postnatal days

**Supplemental Figure S2.** Microbiota diversity. **(A)** Microbial α-diversity (Chao 1 index) and **(B)** microbial β-diversity (Bray-Curtis index) in the feces at PND8 and in the ileal and colonic digesta and feces at PND24 of formula-fed (CTRL, AI, HI) and sow milk-fed (SM) piglets. A: a,b,c non-common letters indicate that piglet groups differed significantly, p < 0.05. A three-way ANOVA was performed testing for main effects of diet, sex, litter, and for interactions between diet and sex, and diet and litter, followed by Tukey *post hoc* test for parametric variables or a Kruskal Wallis test associated with Dunn test for non-parametric variables. B: no difference in β-diversity occurred when only the three formula groups were compared (PERMANOVA p > 0.05).

**Supplemental Figure S3.** Correlation matrix with Benjamini–Hochberg correction between relative abundance of microbial genera and physiological variables in the ileum **(A)** and colon **(B)** of formula-fed (CRTL, AI and HI) piglets at PND24. Only associations with a correlation coefficient |r|> 0.7 and a significant p value < 0.05 are shown, with the strength of those correlations denoted by the intensity of the circle’s colour.

Muc cell per 100µm crypt/villous, density of mucin cells in 100µm of crypt/villous; Muc cell per crypt/villous, density of mucin cells per crypt/villous; Crypt depth, crypt depth in µm ; Villous length, villous length in µm ; sIgA, sIgA concentrations in the ileal content ; TNF_T_PEYER, TNF_LPS_PEYER and TNF_ConA_PEYER, TNFα secretion by Peyer Patch cells in unstimulated condition (T) or stimulated by LPS or ConA; Total SCFA, total SCFA concentration; short chain fatty acid content is expressed in raw value or ’…_prop’ in % of total SCFA content; the variables in capital letters correspond to the expression of genes listed in Supplemental Table S7.

↓ indicates a genera that is significantly correlated with at least an OTU potentially corresponding to AI and/or HI SynCom strains (see Figure 7).

Supplemental Table S1. Nutritional composition of formula and sow milk.

Supplemental Table S2. Percentage of identity between the 16S rRNA gene of each SynCom member and the corresponding OTU sequence, as determined using BLAST.

**Supplemental Table S3.** Differential abundances of OTUs in the ileum, colon and feces at PND24 and the feces at PND8 of formula-fed (CRTL, AI and HI) piglets.

**Supplemental Table S4.** Relative abundances of families and genera in the feces at PND8 and in the ileum, colon and feces at PND24 of formula-fed (CRTL, AI and HI) piglets and sow milk-fed (SM) piglets, and the differential abundance between formula-fed and sow milk-fed groups (log2 fold change).

**Supplemental Table S5**. Cytokine (IL-10 and TNF-α) and secretory IgA (sIgA) production by mononuclear cells extracted from ileal Peyer’s Patches of formula-fed (CTRL, AI, HI) and sow milk-fed (SM) piglets at PND24

**Supplemental Table S6**. Expression of ileal and colonic genes involved in epithelial barrier, immune, tryptophan metabolism and endocrine functions, which were non-significantly different between formula-fed (CTRL, AI, HI) and sow milk-fed (SM) piglets at PND24.

**Supplemental Table S7**. Primers used in real-time PCR.

