## Supplementary figures and images for "Human milk bacteria assembled into functionally distinct synthetic communities in infant formula differently affect intestinal physiology and microbiota in neonatal mini-piglets"

### Supplemental Figure S1.jpg

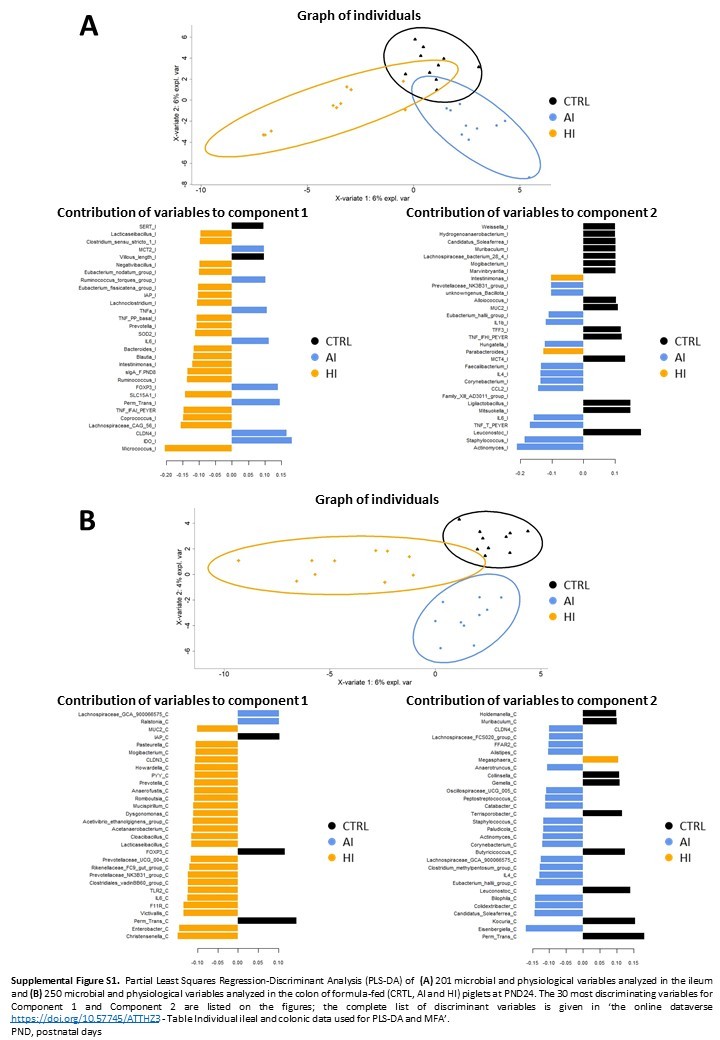

### Supplemental Figure S2.jpg

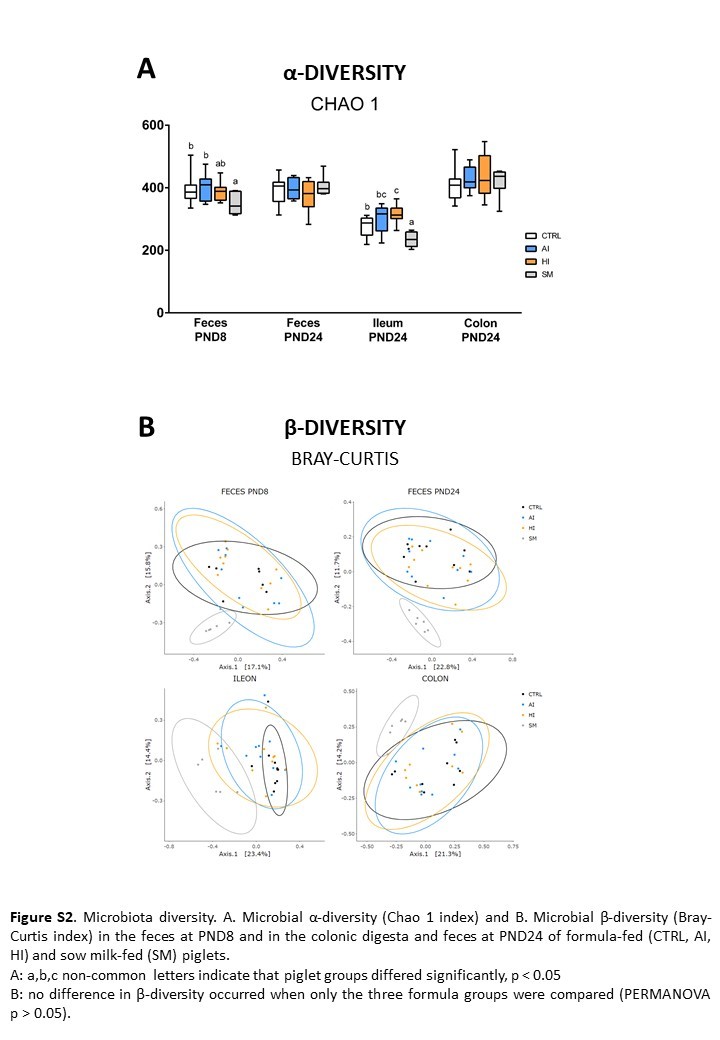

### Supplemental Figure S3.jpg

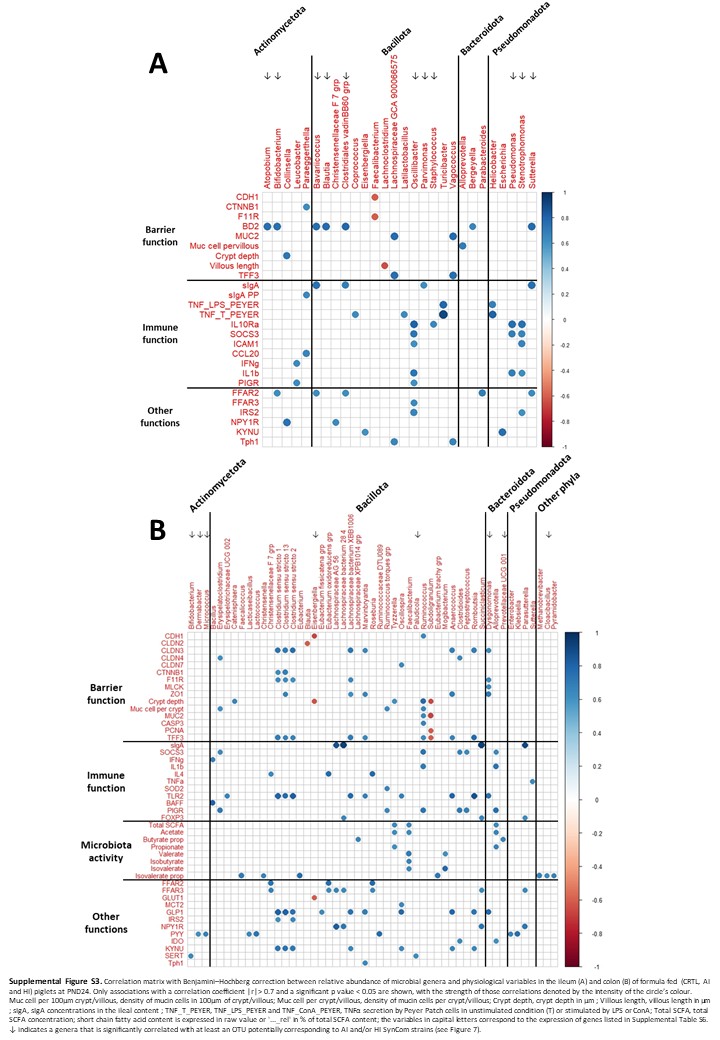
