## Supplemental Table S1 for "Human milk bacteria assembled into functionally distinct synthetic communities in infant formula differently affect intestinal physiology and microbiota in neonatal mini-piglets"

**Supplemental Table S1.** Nutritional composition of formula and sow milk.

| | Formula[1] | Sow milk[2] |
|---|---|---|
| Total powder (g / 100 mL) | 20.00 | |
| Total dry matter (g / 100 mL) | 19.40 | 20.10 |
| Proteins (g / 100 mL) | 3.43 | 5.11 |
| Lipids (g / 100 mL) | 8.42 | 7.24 |
| Carbohydrates (g / 100 mL) | 6.21 | 7.00 |
| Ashes (g / 100 mL) | 1.34 | 0.77 |
| Energy (kcal / 100 mL) | 119.5 | 113.5 |

[1], Trace minerals and vitamins supplied per 100 g of powder: Ca 750 mg, K 600 mg, P 484 mg, Cl 360 mg, Na 200 mg, Mg 32 mg, Fe 13 mg, Zn 12.5 mg, Mn 4.2 mg, Cu 2 mg, Se 39 µg ; vitamin A 640 µg ER, thiamin (B1) 0.45 mg, riboflavine (B2) 1.2 mg, niacin (B3) 5.95 mg, pantothenic acid (B5) 4.1 mg, vitamin B6 0.5 mg, vitamin B8 15 µg, vitamin B9 182 µg EFA, vitamin B12 2.8 µg, vitamin C 75 mg , vitamin D 16 µg, vitamin E 20 mg a-TE, vitamin K 42 µg.

[2], Yucatan milk composition at PND21 (mean value of the milk samples from four sows)
