## Supplemental Table S5 for "Human milk bacteria assembled into functionally distinct synthetic communities in infant formula differently affect intestinal physiology and microbiota in neonatal mini-piglets"

**Supplemental Table S5**. Cytokine (IL-10 and TNF-α) and secretory IgA (sIgA) production by mononuclear cells extracted from ileal Peyer’s Patches of formula-fed (CTRL, AI, HI) and sow milk-fed (SM) piglets at PND24

|  | | **Culture condition** | **CTRL** | | | **AI** | | | **HI** | | | **SM** | **p value** |
| --- | --- | --- | --- | --- | --- | --- | --- | --- | --- | --- | --- | --- | --- |
|  | IL-10^1^ (pg/mL) | Basal | nd | | | nd | | | nd | | | na |  |
|  |  | ConA-stimulated | 329 | ± | 112 | 274 | ± | 95 | 353 | ± | 87 | na | 0.373 |
|  |  | LPS-stimulated | 5.0 | ± | 2.2 | 4.9 | ± | 2.4 | 5.3 | ± | 1.7 | na | 0.751 |
|  | TNFα^1^ (pg/mL) | *Basal* | *1.85* | *±* | *0.46* | *8.96* | *±* | *3.35* | *7.38* | *±* | *2.45* | na | *0.053* |
|  |  | ConA-stimulated | 116 | ± | 35 | 84 | ± | 31 | 104 | ± | 29 | na | 0.665 |
|  |  | LPS-stimulated | 16.4 | ± | 10.7 | 8.4 | ± | 4.0 | 14.6 | ± | 6.4 | na | 0.734 |
|  | sIgA^2^ (pg/mL) | Basal | 179 | ± | 68 | 107 | ± | 28 | 162 | ± | 63 | 47 ± 19 | 0.148 |

^1^ ileal Peyer’s Patch cells were cultured in unstimulated (basal) or stimulated (Concavalin A, Con A or LPS) conditions during 2 days for quantification of cytokine production ; ^2^ ileal Peyer’s Patch cells were cultured during 7 days for quantification of sIgA production

Data are presented as means ± SEM ; na, non-available; nd, not detected as values were under the limit of detection ; italic indicates a tendency with p-values between 0.05 and 0.1.

PND, postnatal days.
